# The transcription factor Chronophage/BCL11A/B promotes intestinal stem cell proliferation and endocrine differentiation

**DOI:** 10.1101/2024.08.05.606739

**Authors:** Emer Aisling King, Eleanor Jacobsen, Nicholas Woolner, Joaquín de Navascués, Owen J Marshall, Jerome Korzelius

## Abstract

Tissue-resident Adult Stem Cells (ASCs) need to continuously adapt their rate of division and differentiation based on their tissue environment. However, the gene regulatory networks that govern these decisions in ASCs and how they respond to challenges such as infection are often not fully understood. We identify a novel role for the transcription factor (TF) Chronophage (Cph) in ISC proliferation and entero-endocrine (EE) cell differentiation. Cph is a Z2H2 zinc TF orthologous to mammalian BCL11A/B that are involved in regulating adult stem cell fate in various contexts. We show here that Cph is expressed in ISCs and EEs in the *Drosophila* intestine. Increased levels of Cph correlates with increased ISC proliferation and EE differentiation. *cph* loss-of-function leads to impaired ISC proliferation. Cph levels are elevated during tumourigenesis as well as in ageing and infection conditions. Knockdown of Cph in a Notch-mutant tumour model reduces tumour size and incidence and extends lifespan. Mechanistically, Cph overexpression leads to an increase in enteroendocrine (EE) cells and DamID DNA-binding and qRT-PCR analysis reveals that Cph directly regulates the levels of key EE regulatory genes such as Prospero (*pros*) and Phyllopod (*phyl*). In addition, Cph directly regulates core cell cycle regulators such as E2F1 as well as the TF Nerfin-1 that controls ISC proliferation and maintenance. Together, these data support a role for Cph in finetuning the balance between differentiation and proliferation during entero-endocrine differentiation.

## Introduction

Adult stem cell (ASC) dysfunction is a major contributor to organismal decline in ageing and a key driver of age-associated diseases such as cancer (Brunet et al., 2023; Kennedy et al., 2014; López-Otín et al., 2023). Tissue-resident ASCs that maintain various tissues throughout the body have unique combinations of transcription factors (TFs) that determine their maintenance, proliferation, and differentiation. Recent years have seen a renewed interest in the idea of using embryonic stem-cell transcription factor combinations such as Oct4, Klf4, Sox2 and Myc (OKSM) (Takahashi and Yamanaka, 2006) to re-program tissues *in vivo* to rejuvenate ASCs and improve tissue function (Lu et al., 2020; Ocampo et al., 2016, 2016). However, these broad-acting reprogramming factors come with an associated risk of unchecked stem/progenitor cell expansion and a risk of teratoma formation (Abad et al., 2013; Parras et al., 2023). Ageing also disrupts the differentiation/proliferation balance in adult tissues (Brunet et al., 2023) such as the intestine and lead to oncogenic transformation (Schwitalla et al., 2013). A more bespoke strategy for tissue rejuvenation would require a combination of ASC-specific transcription factors tailored to its tissue that would maintain the proper balance between stem cell proliferation and differentiation. This highlights the need to understand the network of TFs that maintain ASC function with age in each tissue and leverage this understanding to design novel strategies to prevent age-associated tissue decline.

The intestinal stem cells (ISCs) resident in the adult midgut or intestine of the fruit fly *Drosophila melanogaster* offer a well-characterised system ideal for studying the regulators of ASC proliferation, maintenance and differentiation in the context of ageing (Jasper, 2020). ISCs in the fly intestine divide to self-renew and generate either an absorptive enterocyte (EC) or secretory enteroendocrine (EE) lineage by way of a pre-committed daughter cell (Micchelli and Perrimon, 2005; Ohlstein and Spradling, 2005; Perdigoto et al., 2011; Zeng and Hou, 2015). Age-associated changes in the intestine have been found to contribute to stem/progenitor misdifferentiation and an increase in EE-progenitors (Biteau et al., 2010, 2008; Tauc et al., 2021). Notch signalling is central to the process of ISC differentiation, with high levels of Notch signalling from the ISC to the precursor cell leading to the formation of an enteroblast (EB), an EC precursor cell. Conversely, low levels of Notch signalling leads to the generation of a pre-EE cell and subsequently an EE cell (Ohlstein and Spradling, 2007; Perdigoto et al., 2011). Recent studies also suggest direct differentiation of ISCs to EE’s (Zeng and Hou, 2015). Loss of Notch signalling leads to the generation of neoplastic tumours consisting of ISC-like cells and EE cells (Ohlstein and Spradling, 2007, 2005; Patel et al., 2015; Perdigoto et al., 2011). Active Delta/Notch signalling between the ISC and its daughter results in the daughter adopting the EB-progenitor fate, characterized by high Notch activity. Upon activation by cleavage, the Notch intracellular domain moves into the nucleus and acts as a transcription factor by activating several other target genes that ensure correct differentiation of the EB to the EC-fate.

One of the critical factors that play a role in safeguarding lineage identity is the Wilms Tumour 1 (WT1)-like zinc-finger TF Klumpfuss (Klu) (Yang et al., 1997). In the adult intestine Klu acts downstream of Notch in EBs and loss of Klu results in EB mis-differentiation towards the EE fate (Korzelius et al., 2019). High levels of Klu lead to apoptosis of EBs, which seems to be another role in which Klu controls EB differentiation rates (Reiff et al., 2019). *Klu* ensures EC cell fate in EBs by repressing genes involved in ISC proliferation and EE differentiation. Through the study of a set of high-confidence Klu target-genes, we identified the Z2H2 zinc transcription factor CG9650/Cph as a novel regulator of ISC proliferation and EE differentiation.

CG9650 was recently dubbed *Chronophage* (*cph)* for its role in temporal patterning of the central nervous system (CNS) (Fox et al., 2022). Cph represses Pdm1 in neural stem cells (NSCs) allowing for the transition to Castor-expressing neurons during development of the ventral nerve cord (Fox et al., 2022; Tang et al., 2022). Cph was also identified in a gain-of-function screen of CNS development modulators (McGovern et al., 2003). Alongside this, Cph has been shown to modulate levels of Notch expression in developing cone cells during *Drosophila* eye development (Shalaby et al., 2009). Cph has two mammalian homologs CTIP1 (BCL11A) and CTIP2 (BCL11B) and is thought to be a co-factor for the BAF-Swi/Snf chromatin remodelling complex. Cph/BCL11A has been identified as a Type 2 diabetes risk gene and has been shown to regulate insulin secretion in human islet cells and *Drosophila* insulin-producing cells (Peiris et al., 2018). Cph has also been identified as a factor involved in modulating insulin secretion (Park et al., 2014), demonstrating a level of functional conservation between Cph and its human homologs.

This work demonstrates a key role for Cph in controlling ISC proliferation and EE differentiation. Cph is expressed in a sub-population of ISCs and EEs under homeostatic conditions but is strongly induced in situations of ISC overproliferation such as in Notch mutant tumours and in the aged intestine. Loss of Cph leads to defects in ISC proliferation, whereas Cph overexpression is sufficient to induce partial EE differentiation in stem/progenitor cells. Mechanistically, Cph controls ISC proliferation and EE differentiation through binding of key EE-differentiation factors such as the EE master regulator Prospero (Pros) (Guo et al., 2022a; Li et al., 2017) and Phyllopod (Phyl) (Yin and Xi, 2018). Furthermore, Cph controls ISC proliferation by regulation of core cell cycle regulators such as E2F1 and Cyclin E as well the TF Nerfin-1, which we found to control ISC maintenance and proliferation in response to damage. Altogether, these results uncover a central role for Cph in controlling the balance between proliferation and differentiation in adult stem cells.

## Results

### Cph is expressed in the stem and entero-endocrine cells of the adult intestine

From our analysis of Klu high-confidence target genes that were 1. bound directly by Klu 2. upregulated by klu knockdown and 3. downregulated by Klu-overexpression we identified the zinc-finger TF CG9650/Cph. We first reviewed the scRNA-Seq atlas of the *Drosophila* adult intestine, generated by the Perrimon lab to query Cph expression in this tissue (Hung et al., 2020). This showed Cph to be expressed in ISCs and EEs. Furthermore, scRNAseq analysis of EE subtypes showed Cph to be expressed in most EE subtypes defined by Guo et al. (X. Guo et al., 2019). To confirm the expression pattern of Cph we used a *cph-YFP^CPTI1741^*protein trap line from the Cambridge protein Trap Initiative (CPTI, (Lowe et al., 2014). Cph was found to be expressed in neural stem cells and neurons in the larval central nervous system (Fox et al., 2022). We therefore decided to stain this line for stem cell and entero-endocrine cell markers for the intestine. We first stained the *cph-YFP* protein trap line with the EE marker Prospero (Pros) (red) (Fig 1a-a’’). We observed Cph expression in a subset of Pros+ EE cells (Figure 1a-a’’, arrowheads), but absent from others (Fig1a-a’’, arrows). To examine the overlap in Cph expression with differentiated EE subtype markers such as the Class II EE marker, the gut hormone Tachykinin (Tk), we used a gut-specific *Tk_gut_-Gal4* line (Song et al., 2014) (Supplementary Figure 1). Cph appeared to be expressed in all Tk+ cells in the R5 region however it is not exclusively expressed in Tk+ cells. Hence, Cph expression is expressed in differentiated TK+ EE cells. To address expression of Cph in ISCs, the *cph-YFP* line was crossed to the stem cell-specific reporter *Delta-lacZ* (*Dl-lacZ*) (Figure 1 b-b’’). Cph expression overlapped with *Dl-lacZ* in most cases (Figure 1b-b’’, arrowheads), but we also identified cases of *Dl-lacZ*-positive ISCs without detectable *cph-YFP* expression (Figure 1b-b’’, arrows). This showed that Cph is expressed in a subset of ISCs. To further examine the expression of Cph in the stem-progenitor compartment, we used the stem-progenitor specific Gal4 driver *escargot* (Esg), which expresses exclusively in ISCs and EB precursor cells. (Supplementary Figure 1a-a’’’). Cph was seen to be expressed in a subset of Esg^+^ cells. To investigate whether Cph was expressed in both ISCs and EBs we crossed the *cph-YFP* line to *Su(H)GBE-LacZ*, a Notch-responsive element which is active in Notch-positive EB progenitor cells (Furriols and Bray, 2000; Micchelli and Perrimon, 2005; Ohlstein and Spradling, 2007). We found Cph was not expressed in EBs (Figure 1c-c’’ arrows) and was often found in the small, diploid cells adjacent to the EB: the presumptive ISC mother cell (Figure 1c-c’’, arrowheads). Finally, we performed pseudo-time analysis of *cph* mRNA expression from the integrated intestinal scRNA-seq data sets of the fly gut atlas (Hung et al., 2020) and the fly cell atlas (Li et al., 2022) to examine *cph* expression along the differentiation trajectory into either the EC or EE lineage (Figure 1d). This shows that *cph* expression increases along the EE differentiation trajectory but decreases in the EC differentiation axis (Figure 1e). This further supports our observations with reporter lines and suggests functions for Cph in the ISC and EE cells of the gut.

**Figure 1.**
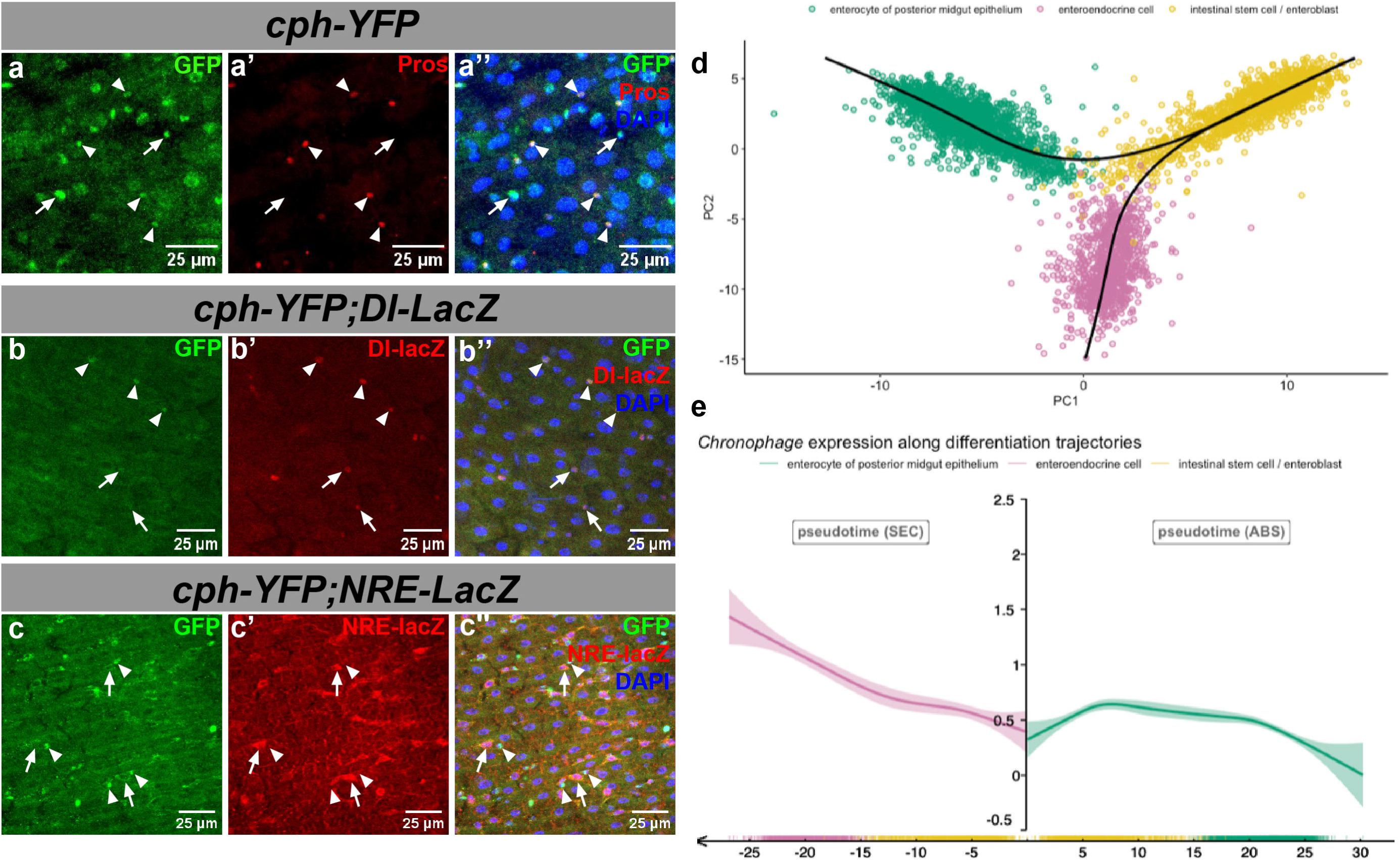
The transcription factor CG9650/Cph is expressed in ISCs and a subset of EEs in the adult *Drosophila* intestine. **a-a’’** The *cph-YFP^CPTI-1740^* protein trap line (*cph-YFP*, green) was dissected and stained with EE marker Prospero (Pros, red). *cph-YFP* is expressed in a subset of Pros-positive EE cells (arrowheads). We also see Cph is expressed in other small diploid cells that are not Pros^+^; most likely ISCs (arrows). *n*=26 animals across 3 biological repeats. **b-b’’** *cph-YFP^CPTI-1740^*was crossed to the ISC-specific *Delta-LacZ* reporter and stained with anti-beta-galactosidase antibody (red). *cph-YFP* is expressed in a subset of Delta^+^ ISCs (arrowheads), arrows point to Dl^+^ Cph^-^ cells demonstrating Cph is not expressed in all Dl^+^ ISCs**. c-c’’’** The *cph-YFP^CPTI-1740^* line was crossed to the Notch activity reporter *Su(H)GBE-LacZ* which marks EBs. Staining with anti-beta-galactosidase (red) we see that *cph-YFP* is not expressed in EB’s (arrows) but in adjacent cells: the presumptive ISCs (arrowheads). **d-e** Pseudo-time analysis was carried out of the expression of *cph* mRNA using data from the single-cell Gut Atlas and Fly Cell atlas (see Methods). **d** The differentiation trajectories for both EE and EC lineages from the ISC. **e** *cph* expression is seen to be increased in the EE lineage and decreased along the EC lineage. *cph* expression also appears to increase in the EE lineage with time.

Based on the above expression analysis, Cph is expressed in a subset of ISCs and EEs. We therefore wanted to confirm that Cph expression marks a stem cell population that would be able to establish clones of cells in the midgut in a similar manner as established ISC markers Delta and Esg. To this end, we used a *cph-Gal4* reporter line (*1151-Gal4*). To verify that this driver reflects the expression of *cph* in the midgut, we crossed this to a *UAS-Stinger-GFP; mira-His2A-mCherry* line. Stinger-GFP is a widely used nuclear-localized GFP (Barolo et al., 2000) and *miranda* (*mira*) is a well reported stem/progenitor cell specific gene (Bardin et al., 2010). This showed co-localisation of *cph-Gal4*-driven *UAS-Stinger-GFP* and *mira-His2A-mCherry* (red, arrowheads) expressing cells (Figure 2 a-a’’). We did not observe any Cph expression in the large polyploid absorptive enterocyte population (Figure 2 a-a’’ asterix), confirming that Cph is expressed in a subset of ISC and EE cells. Then we crossed this *cph-Gal4* reporter line with a line carrying a FlipOut cassette (*tub-Gal80^ts^-inducible UAS-GFP; UAS-FLP, Act5C>CD2STOP>Gal4*) that will lineage trace *cph*-expressing cells and their daughter cells using an inducible *Actin5C-Gal4* cassette that becomes activated by UAS-Flp activation by *cph-Gal4* (hereafter named *cph> F/O* (Jiang et al., 2009). Clones were allowed to develop for 14 days before being dissected (Figure 2 b-e). Guts were stained with the EE marker Pros and the EC marker Pdm1 (Figure 2 f-g’’). No clones were generated in the posterior midgut of the negative FlipOut control without Gal4, although some single EC clones were seen in the anterior midgut of 2/9 animals analysed (Figure 2 b-c). In contrast, *cph> F/O* clones were large and contained both Pros^+^ EEs and Pdm1^+^ ECs within the labelled area (Figure 2 f-g’’). Hence, *cph* is expressed in ISCs that can actively proliferate and differentiate into either EEs or ECs.

**Figure 2.**
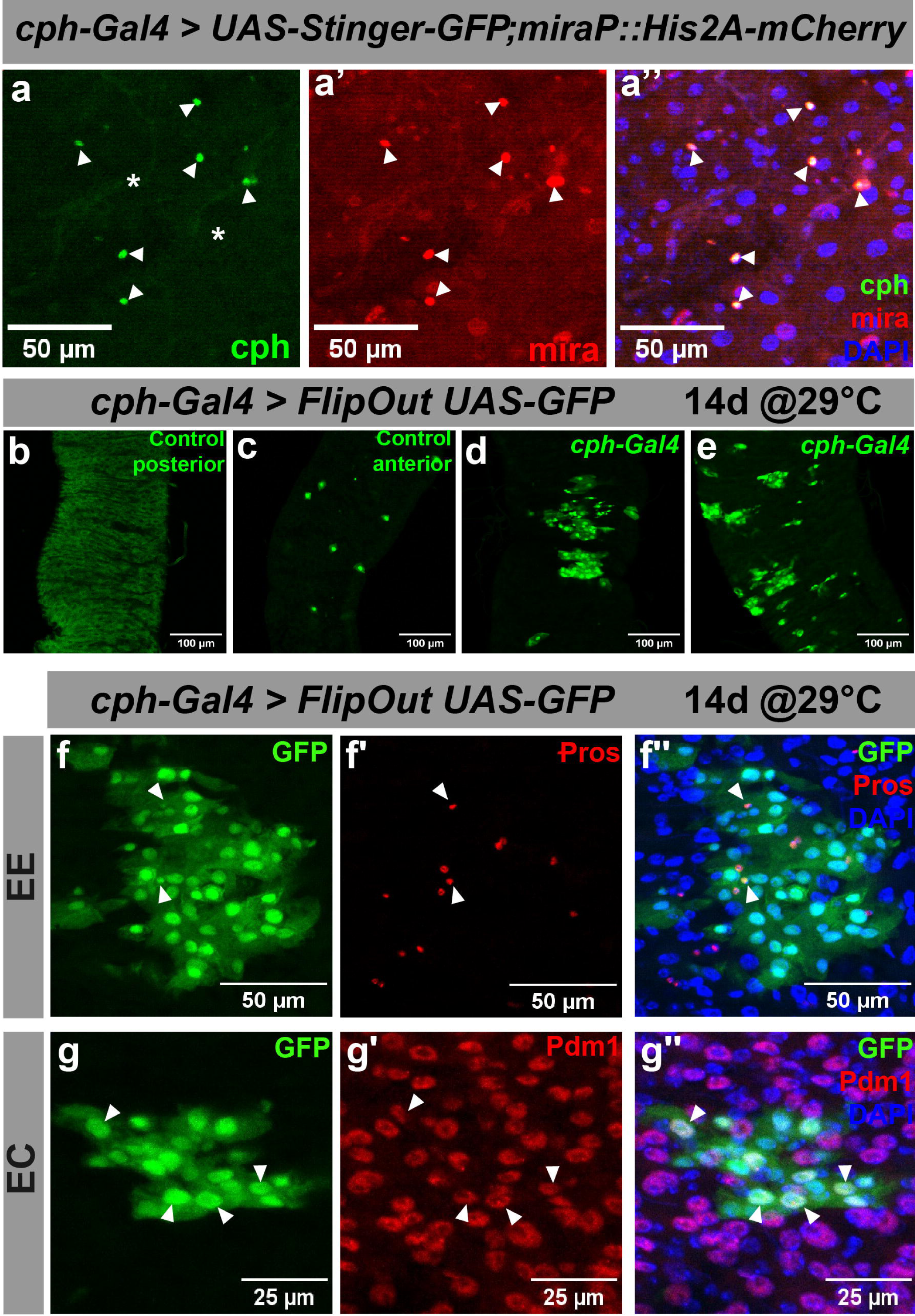
Cph-expressing ISCs are bipotent and differentiate into both EEs and ECs. **a-a’’** The *1151-Gal4* (*cph-Gal4)* line was crossed to a *UAS-Stinger-GFP; mira-His2A-mCherry* reporter line, marking *cph*^+^ cells in GFP (green) and *mira*^+^ progenitor cells in red. Cph is expressed in *mira*^+^ progenitor cells (arrowheads) but absent in polyploid ECs (asterisk). **b-e** The *cph-Gal4* line was crossed to a temperature-inducible FlipOut (F/O) cassette (*w; tub-Gal80ts, UAS-GFP/CyO,wg-LacZ; UAS-FLP, Act5C>CD2STOP>Gal4/TM6B). cph-Gal4>F/O* clones were induced at 29°C and dissected 14 days post clonal induction. *cph-Gal4*>*F/O* guts generated clones (d-e) whereas clones were virtually absent from the posterior region of the negative control guts (b), with some sporadic 1-cell clones in the anterior (c). **f-g** Guts were stained with either the EE marker Pros (red) or the EC marker Pdm1 (red) to assess EE and EC-differentiation respectively. **f-f’’** *Cph-Gal4>F/O* clones containing Pros^+^ EE cells (arrowheads). **g-g’’** *Cph-Gal4>F/O* clones containing Pdm1^+^ EC cells (arrowheads).

### Cph expression is increased during ageing and infection

The fly intestine is a highly dynamic organ and its size and stem cell number is affected by ageing, food availability, mating and different stressors such as enteric infection (Amcheslavsky et al., 2009; Buchon et al., 2010, 2009; Hudry et al., 2016; Jiang et al., 2009; O’Brien et al., 2011).Given the expression pattern of Cph in a sub-set of ISCs, we investigated whether the Cph expression pattern would be altered under stress. To induce stress, we infected *cph-YFP* animals with the pathogenic bacteria *Erwinia carotovora carotovora* (*Ecc15*) (see Methods). 24 hours post-infection animals were dissected along with uninfected controls. Guts were stained for the mitotic marker phosphorylated Histone H3-Ser10 (pH3) to allow detection of increased ISC proliferation. Levels of Cph-YFP fluorescence intensity were taken as a proxy for Cph expression as outlined in the Methods section. We found levels of fluorescence intensity to be significantly higher in *Ecc15*-infected and 5% sucrose fed animals compared to the animals maintained on a regular diet (Figure 3a-d). We did not however observe a significant difference in fluorescence intensity between the 5% sucrose fed control and the infected guts. In our experience, 5% sucrose feeding leads to enhanced ISC proliferation, even without *Ecc15* bacteria added to the sucrose-mix. Nevertheless, comparing these results to the regular diet control, we conclude that Cph expression is induced in these conditions of increased ISC proliferation.

**Figure 3.**
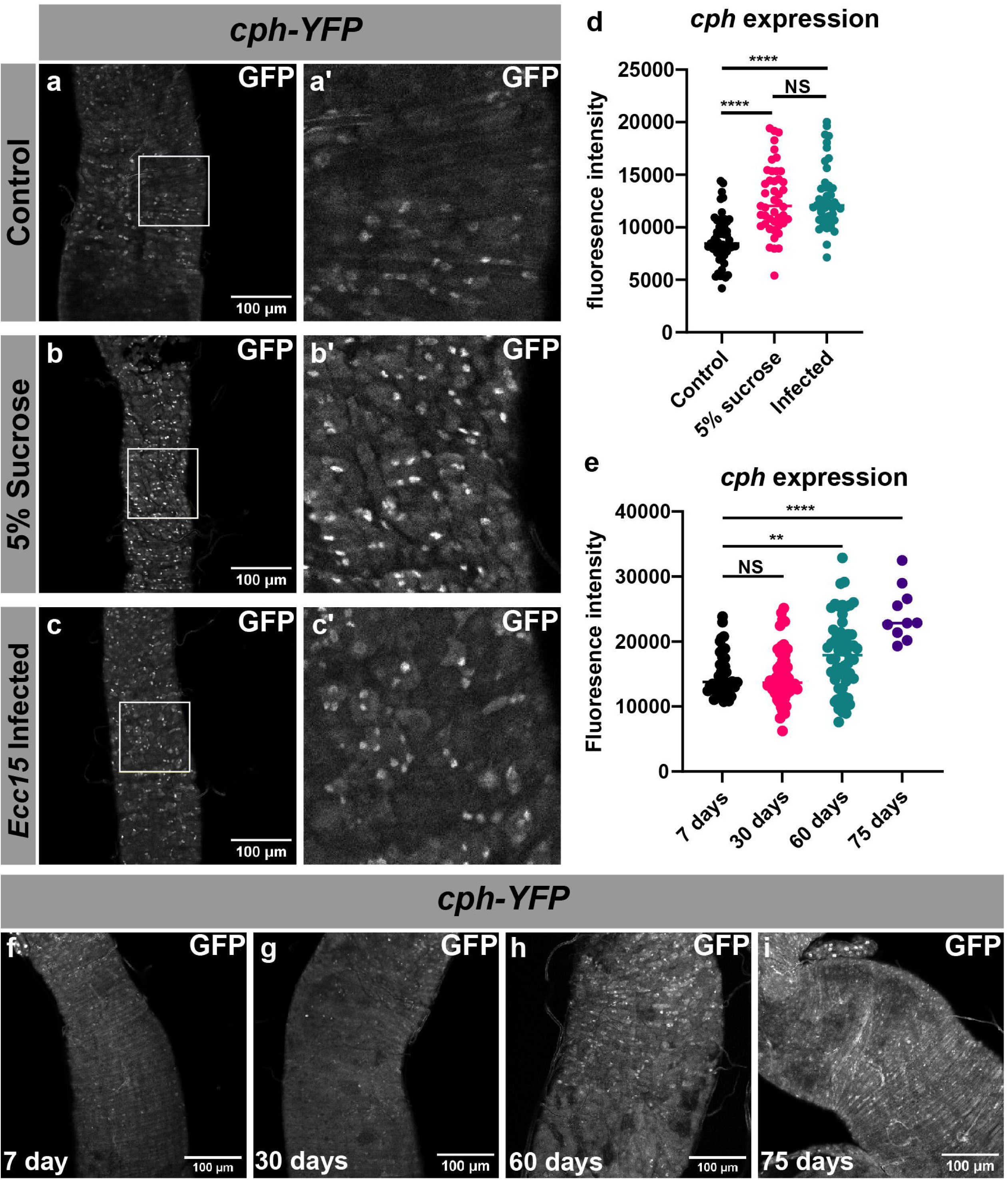
Cph expression is increased during ageing and infection. **a-c’** The *cph-YFP^CPTI-1740^* line was infected with pathogenic *Ecc15* bacteria to induce ISC proliferation and gut turnover (see Methods). Cph-YFP expression is low in control animals (**a-a’**) but is increased in 5% sucrose fed (**b-b’**) as well as in *Ecc15* infected animals (**c-c’**). **d** Quantification of *cph-YFP^CPTI-1740^* fluorescence. Scatter plot displays fluorescence intensity values for each gut and condition. This shows an increase in fluorescence intensity of infected (*p*=3.3e-11) and 5% sucrose fed *cph-YFP* flies (*p*=2.8e-9) compared to control. Control *n*=53, 5% sucrose *n*=45, *Ecc15 n*=44. **e-i** Animals were collected, reared at 25°C and dissected at 7, 30, 60 and 75 days of age. Quantification in **e** shows a significant increase in fluorescence intensity from the 7-day to the 60- and 75-day old flies (7 vs 60 days, *p*=0.0023; 7 vs 75 days, *p*=3.6e-9). 7-day *n*=38, 30-day *n*=51, 60-day *n*=60, 75-day *n*=10. Significance is represented as *, **, *** and **** for *p* < 0.05, 0.01, 0.001 and 0.0001, respectively, comparing fluorescence intensity at 75 days with all other ages (7 vs 75 days, *p*=3.6e-9; 30 vs 75 days, *p*=1.3e-8; 60 vs 75 days, *p*=0.001).

Alongside infection, age is also associated with increased dysbiosis and an increase in ISC proliferation, and more so in egg-laying females than males (Biteau et al., 2010; Buchon et al., 2009; Jasper, 2020; Regan et al., 2016). Having observed a correlation between increased proliferation and expression of Cph, we decided to investigate the expression pattern of Cph in aged *cph-YFP* females. Animals were dissected at 7, 30, 60, and 75 days of age. Cph-YFP levels were found to be significantly increased at the 75-day timepoint compared to all younger timepoints (Figure 2 e-i). No statistically significant difference was found between young (7-day) and middle-aged (30-day flies), whereas the 60-day and 75-day females had significantly more Cph expression (Figure 2e). This is in line with scRNA-seq data obtained from the Aging Fly Cell Atlas for both the EE and ISC/EB cluster (Supplementary Figure 2a-b) (Lu et al., 2023). In concordance with earlier results we also observed a significant increase in Pros^+^ EE cells in older females (Supplementary Figure 2c-d) (Choi et al., 2008; He et al., 2018; Tauc et al., 2021). Alongside this we also noted a timewise significant increase gut cellularity measured by the total number of DAPI^+^ cells present within each gut (Supplementary Figure 2e-f). Altogether, these data show that Cph is induced in conditions of increased ISC activity and gut cell turnover such as infection and ageing.

### Cph loss impairs ISC proliferation

Our data suggest a role for Cph in ISCs, as it is expressed in these cells (Figure 1) and it is also induced in the context of increased ISC activity (Figure 2). Hence, we investigated whether loss of *cph* would impact ISC proliferation. We employed the Mosaic Analysis with a Repressible Cell Marker (MARCM) system to generate GFP-marked *cph* homozygous null clones (Fig 4 a’-f’) (Lee and Luo, 1999). The experiment were carried out with the independent alleles *cph^A^* and *cph^B32^,* both null (Manfred Frasch, personal communication). Clones were induced by heat-shock at 37°C and allowed to grow for 3 and 7 days. Quantification of clone size revealed both *cph^A^* and *cph^B32^* null clones had significantly fewer cells per clone compared to control (FRT19A) clones at both the 3-day (Supplementary Figure 3b-b’’) and 7-day timepoint (Figure 4 a-f’, quantification in Figure 4 g). This suggests that loss of *cph* negatively impacts ISC proliferation. In addition, *cph^A^* and *cph^B32^* animals had significantly fewer clones in their gut overall (Figure 4 h). This suggests that apart from proliferation and differentiation, loss of *cph* in clones might also impact cell survival and ISC maintenance. It should be noted that Pros^+^ EEs were still present in some *cph^A^* and *cph^B32^* MARCM null clones (Supplementary Figure 3 b-b’’). This suggests that EE differentiation is not significantly affected by loss of Cph.

**Figure 4.**
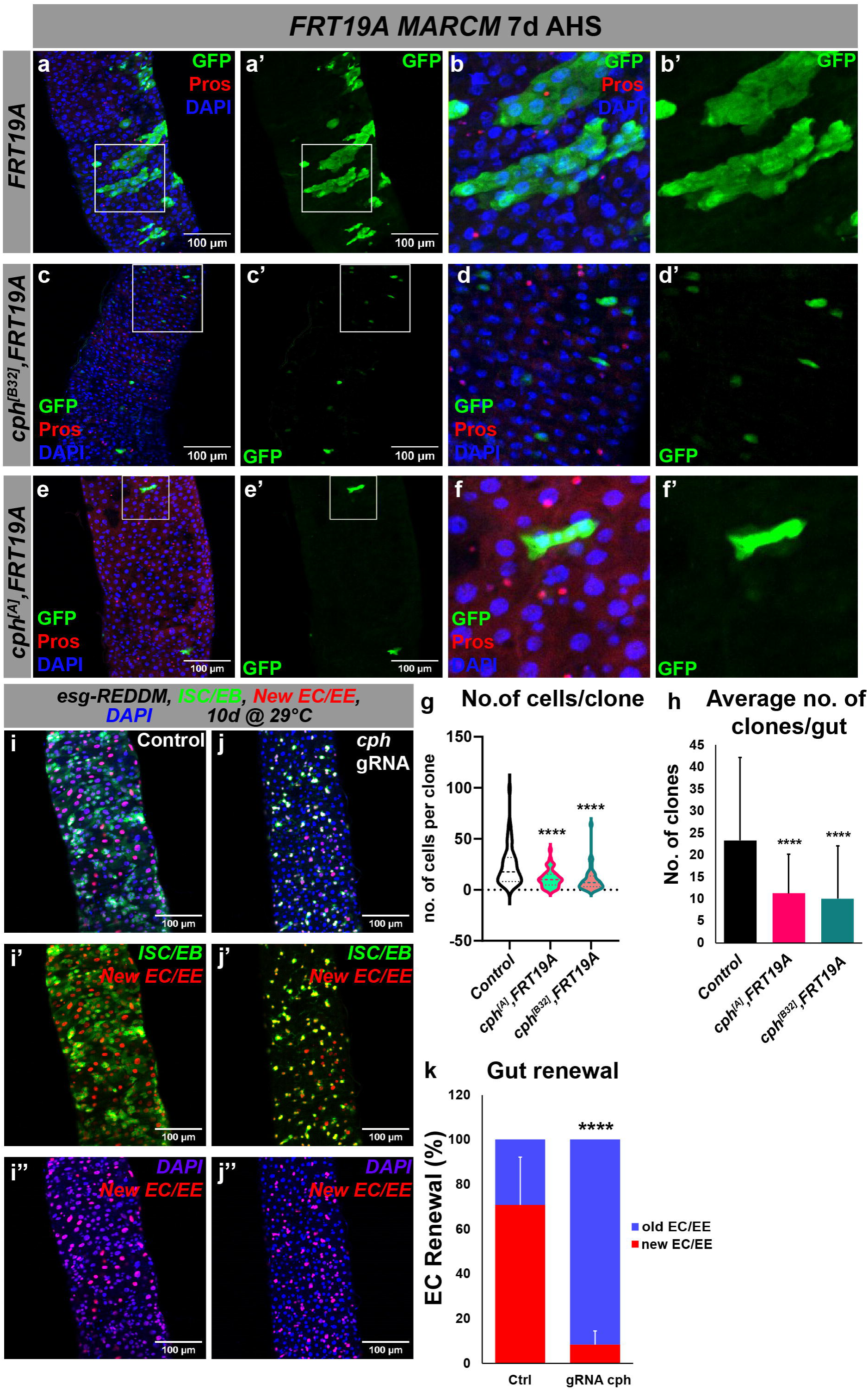
Loss of Cph leads to impaired ISC proliferation. **a-f’** MARCM null clones for *cph* fail to proliferate. *FRT19A* MARCM-ready flies were crossed to either *FRT19A, cph^[A]^ FRT19A* or *cph^[B32]^ FRT19A*. Appropriate progeny were collected and heat-shocked for 45 minutes at 37°C to induce recombination. Progeny were then placed in 25°C incubators for 7 days before being dissected and immunostained for Prospero (red). AHS stands for after heat shock. FRT19A control clones are large multicellular clones 7 days after induction (**a-b’**), whereas both *cph^[A]^* and *cph^[B32^* are mostly single-cell clones at this timepoint, with little differentiation (**c-f’**) **g** Quantification of total number of cells per clone for the abovementioned genotypes. Average clone size was significantly decreased in the *cph* null clones compared to WT (*p*=2.9e-9 and *p*=3.2e-12 for *cph^A^* and *cph^B32^*, respectively). Data compiled from three biological repeats with the following n-values: FRT19A (control) *n*= 52, *cph^A^ FRT19A n*= 42*, cph^B32^ FRT19A n*=37. **h** The total number of *cph* null clones per gut decreased significantly compared to the control (*p*=1.1e-4 and *p*=1.2e-4 for *cph^A^*and *cph^B32^*, respectively). **i-k** REDDM gut turnover analysis of midguts with *cph* CRISPR/Cas9-mediated loss-of-function. *esg-REDDM > UAS-Cas9* control and *esg-REDDM > UAS-Cas9, UAS-cph^gRNA^* were placed at 29°C for 10 days to induce REDDM, Cas9 and gRNA before analysis. *cph* CRISPR flies had significantly fewer new EC/EE cells (red/blue) compared to the *esg-REDDM > UAS-Cas9* control. There also appears to be far less ISC/EB proliferation in *cph* CRISPR/Cas9 guts. **k** Quantification of new EC/EE cells in the *cph* CRISPR/Cas9 gut, 10 days at 29°C (*p*= 2.1e-29). Results are based on three independent repeats with the following *n*-values; *esg-REDDM > UAS-Cas9* control *n*= 42, *esg-REDDM > UAS-Cas9, UAS-cph^gRNA^ n*= 78.

To further address a role for Cph in ISC proliferation and cell turnover in the intestine, we used the REDDM system (Antonello et al., 2015) combined with *UAS-Cas9* and crossed this to a line containing a *UAS*-driven *cph* gRNA targeting *cph* to compromise its function by inducible, CRISPR/Cas9-mediated mutation (Port et al., 2020). The REDDM system is based on differential labelling of old and newborn cells to determine cell turnover. To differentiate between newborn and old cells, fluorophores with different half-lives are used. *UAS-CD8-GFP* has a short half-life and exclusively labels stem/progenitor (ISC/EB) cells as it is driven by *esg-Gal4* activity. It is lost from cells such as ECs and EEs once *esg-Gal4* is no longer active. *UAS-H2B-RFP* on the other hand is a stable protein that carries over from the stem/progenitor compartment to differentiated ECs and EEs. Once the system is induced, ISC/EB pairs are tagged with both GFP and RFP (green and red), old EC/EE cells will be marked with the nuclear DNA marker DAPI (blue only) and newborn EC/EE cells are labelled with H2B-RFP (red+blue). Therefore, the ratio of new: old cells (blue VS red+blue) was used as a metric for cell turnover. Crosses were set up and maintained at 18°C before being shifted to the permissive temperature of 29°C for 3 and 10 days to induce the system and *cph* gRNA expression. We found that in *esg-REDDM >UAS-Cas9, UAS-cph-gRNA* guts there is a significant reduction in cell turnover compared to control (*esg-REDDM >UAS-Cas9*) at both the 3-day (Supplementary Figure 3a) and the 10-day timepoints (Figure 4 i-k). We quantified cellular turnover by counting the fraction of new VS old ECs in the above genotypes. Whereas 80% of ECs are quantified as “new” (DAPI+RFP) in control *esg-REDDM >UAS-Cas9* guts after 10 days, *esg-REDDM >UAS-Cas9, UAS-cph gRNA* guts have only around 10% new ECs in the same timeframe (Figure 4 k). Hence, we conclude that Cph is required in ISCs for their proliferation and long-term maintenance.

### Cph is induced in Notch loss of function tumours and is necessary for their expansion

Our findings thus far suggest that Cph is involved in ISC proliferation and maintenance. Our previous work on Klu showed that Cph is directly repressed by the Notch-target Klumpfuss (Klu) which is highly expressed in EBs (Korzelius et al., 2019). Based on these findings and earlier studies suggesting an interaction with the Notch pathway in the developing nervous system (McGovern et al., 2003; Shalaby et al., 2009), we decided to explore the potential interaction between Cph and Notch signalling. The Notch signalling pathway plays a central role in ISC differentiation. Notch knockdown in ISCs and EBs results in neoplastic ISC-like tumours forming. In the absence of Notch, EC differentiation cannot take place and this leads to an excess of EE and pre-EE cells expressing high levels of Pros within the tumours. (Micchelli and Perrimon, 2005; Ohlstein and Spradling, 2005; Patel et al., 2015; Perdigoto et al., 2011). As we found the expression of Cph restricted to ISC and EE cells, but not in ECs, we wanted to analyse the expression pattern of Cph in *Notch*-RNAi induced tumours. To this end, we generated *cph-YFP;UAS-Notch^RNAi^* animals and crossed them to an *esg-Gal4^ts^,UAS-mCherry* driver. The animals were placed at 29°C for 7 days before being dissected and stained with the EE marker Prospero. Cph was found to be widely expressed in the tumour tissue and co-localized with several ISC and EE markers within the tumours: Cph was co-localized with Esg (red, area outlined in Figure 5 b’) and it was also expressed in the precursor EE (pre-EE) type cells (Esg+ and Pros^+^, red and white) as well as in in more differentiated EE cells (Pros only, white) (Figure 5 b-b’’’, arrowhead). Cph-YFP also partially overlapped with the ISC-marker Delta that is highly expressed in Notch tumours (Supplementary Figure 4 a-a’’) Thus, it seems that Cph expression is dramatically induced upon Notch loss of function. To quantify this increase in Cph levels, *esg-Gal4^TS^>Notch^RNAi^*flies were shifted to 29°C to induce tumours, RNA was extracted (see Methods) and used for qRT-PCR analysis to assess the levels of Cph expression. There was a 15-fold increase in *cph* mRNA expression in the *esg-Gal4^TS^>Notch^RNAi^*samples compared to control (*esg-Gal4^TS^>*) (Figure 5c). We conclude that Cph is highly induced in *Notch* loss-of-function tumours.

**Figure 5.**
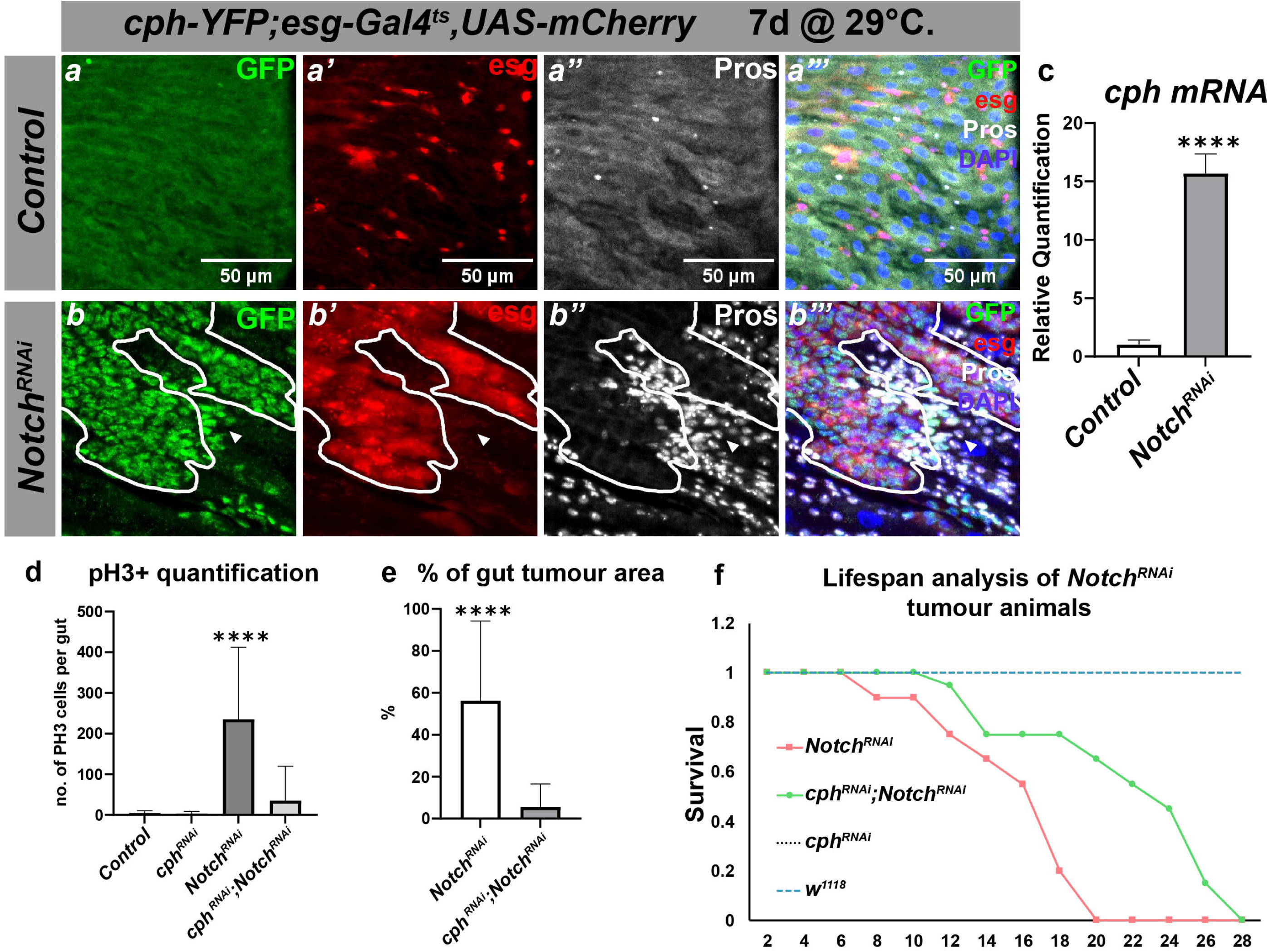
Cph expression is increased in Notch loss-of-function tumours and Cph loss reduces tumour burden. **a-b’’’** *cph-YFP^CPTI-1740^;esg-Gal4^ts^,UAS-mCherry >UAS-Notch^RNAi^* shows Cph-YFP expression in *Notch-RNAi* induced tumours. Animals were dissected after 7 days of induction at 29°C and stained with the EE marker Prospero (Pros, greyscale). **a-a’’’** shows the expression of *Cph* in *cph-YFP^CPTI-1740^; esg-Gal4^ts^ > UAS-mCherry* control guts. **b-b’’’** Cph expression in *esg-Gal4^ts^ > UAS-mCherry, UAS-Notch^RNAi^* guts. Note the distinct overlap in expression of *Cph* and the ISC/EB-marker *escargot* (*esg*) in the tumours outlined in white. There are also tumour cells that are Pros^+^ and Cph^+^ (arrowhead). This experiment was carried out three independent times (compiled n-values are as follows*: esg-Gal4^ts^ > UAS-mCherry n*=27*, cph-YFP^CPTI-1740^;esg-Gal4^ts^ >UAS-Notch^RNAi^ n=*80). **c** qRT-PCR of *cph* mRNA in *esg-Gal4^ts^ >UAS-Notch^RNAi^*14 days after induction alongside *esg-Gal4^ts^ > UAS-mCherry* control flies. *n*=15 guts were dissected from each genotype and used for total RNA extraction and cDNA generation. Data show a 34-fold increase in *cph* expression in the *esg-Gal4^ts^> UAS-Notch*^RNAi^ flies compared to *esg-Gal4^ts^ > UAS-mCherry* control flies (*p*=3.7e-4). **d** *cph* knockdown decreases number of mitotic cells in *Notch ^RNAi^* guts. Quantification of mitosis in *esg-Gal4^ts^ > UAS-mCherry control*, *cph^RNAi^, Notch*^RNAi^ and *cph^RNAi^;Notch*^RNAi^ flies. Females of the appropriate genotype were collected, induced at 29°C and dissected after 7 days. Guts were stained with anti-pH3S10 and counted along the length of the intestine. There was a significant decrease in the numbers of pH3^+^ cells in the *cph^RNAi^;Notch*^RNAi^ compared to the *Notch*^RNAi^ guts (*p*=2.0e-5). The bar chart was compiled from two biological repeats. *cph^RNAi^;Notch*^RNAi^ *n*=51, *Notch^RNAi^ n*=23, *Cph^RNAi^ n*= 14, *esg-Gal4^ts^ > UAS-mCherry n*= 15. **e** Percentage of the portion of gut taken up by tumours in *cph^RNAi^;Notch*^RNAi^ guts is significantly reduced compared to *Notch^RNAi^*(*p*1.3e-6).**f** Loss of *cph* significantly extends lifespan of *Notch*^RNAi^ tumour-bearing flies (*p*= 4.2e-5). Data in Kaplan-Meier curve is representative of 3 independent repeats using two independent *cph* RNAi lines (see Supplementary Figure 4).

To further investigate the dependency of Notch tumours on Cph expression, we generated *Notch^RNAi^* tumours in the presence of absence of a *cph^RNAi^* construct. Animals were placed at 29°C for 7 days before being dissected and stained with the mitotic marker pH3S10 and the EE marker Pros. Total number of pH3^+^ cells were counted as an indicator for ISC proliferation. *esg-Gal4^TS^>cph^RNAi^;Notch^RNAi^* guts had significantly fewer PH3^+^ cells compared to the *esg-Gal4^TS^> Notch^RNAi^* control (Figure 5 d). This indicates that loss of Cph in *Notch^RNAi^*tumours repressed ISC proliferation. The guts were also quantified for tumour growth (see Methods for more detailed description of analysis). There was a 1.9-fold increase in tumour growth in the *esg-Gal4^TS^> Notch^RNAi^* control compared to *esg-Gal4^TS^>Cph^RNAi^;Notch^RNAi^*animals (Results are based on two independent repeats, n-values are as follows *Gal4^TS^>cph^RNAi^;Notch^RNAi^* n=42, *esg-Gal4^TS^> Notch^RNAi^ n=22).* Tumour growth was also slowed significantly in *esg-Gal4^TS^>cph^RNAi^;Notch^RNAi^*compared to control (*esg-Gal4^TS^> Notch^RNAi^*) as assessed by total area of tumour/gut (Figure 5 e, Supplementary Figure 4b-b’’). Finally, a survival assay was carried out to see if reducing Cph-levels in Notch tumours could extend lifespan by reducing tumour burden (see Methods). *esg-Gal4^TS^>Cph^RNAi^;Notch^RNAi^*animals had a significant lifespan extension, surviving on average 8.333 days longer than *esg-Gal4^TS^> Notch^RNAi^* control animals (Figure 5 f). Results were confirmed by another independent *Cph^RNAi^* line (Supplementary Figure 4c). From these results we conclude that loss of Cph in *Notch^RNAi^* induced tumours has a tumour-suppressive role, supporting a role for Cph as a positive regulator of ISC proliferation.

### Cph is highly induced in EE-progenitor tumours

Our data support a critical role for Cph in the positive regulation of ISC proliferation proliferation both in homeostasis and tumour formation. We find that Cph is highly induced in *Notch^RNAi^* tumours that consist mainly of ISC/EE-progenitors (Patel et al., 2015). To further explore the role of Cph in EE progenitor tumours, we employed another tumour model that uses knockdown of the transcriptional repressor *tramtrack* (*ttk*). Tramtrack is a transcriptional repressor that has been dubbed a master repressor of EE differentiation (Harrison and Travers, 1990; Wang et al., 2015; Wang and Xi, 2015). Ttk directly targets and represses the proneural gene *scute* (*sc*) (Yin and Xi, 2018). The transient expression of Scute leads to EE differentiation, promoting asymmetric cell division of ISCs to give rise to a pair of EE cells (Bardin et al., 2010; Chen et al., 2018; Li et al., 2017). Ttk activity is controlled by an E3-ligase destruction complex with the E3-ligase Sina and its regulatory adapter protein Phyl. Low Notch signalling results in the upregulation of Phyl, subsequent destruction of Ttk and de-repression of Scute, triggering EE differentiation (Yin and Xi, 2018). The precise regulation of Scute in ISC proliferation and EE-differentiation is of key importance, as the loss of *ttk* in ISCs leads to uninhibited EE progenitor expansion. Loss of *ttk* leads to formation of neuroendocrine-like tumours (NET) in the gut that share many molecular similarities with NSC in the developing fly brain (Berger et al., 2012; Li et al., 2020; Wang et al., 2015; Wang and Xi, 2015). *ttk*-depleted ISCs switch to a neural stem cell-like program in which the neuroblast stem cell determinant Deadpan (Dpn) drives enhanced self-renewal, and the pan-neural differentiation factor Sequoia (Seq) promotes EE differentiation (Li et al., 2020). Loss of *ttk* is even sufficient to drive transdifferentiation of polyploid ECs through de-repression of Pros in these cells (Guo et al., 2024).

To assess whether Cph is upregulated in *ttk^RNAi^*-induced NET-like tumours, we combined the *cph-YFP* reporter with *UAS-ttk^RNAi^*, crossed this to an *esg-Gal4^TS^* driver line and induced at 29°C for 7 days. Cph-YFP expression is highly expressed in the *ttk*-depleted tumours (Supplementary Figure 5d-e’’’). To confirm this induction of *cph*, *esg-Gal4^TS^>ttk^RNAi^*samples were dissected after 7 days and used for RNA extraction and assayed using qRT-PCR and found to be increased on average 29-fold (Supplementary Figure 5a-c). Based on these results, we conclude that Cph is highly induced in the NET-like state of excess neuro-endocrine differentiation triggered by loss of Ttk.

### Cph modulates EE fate determination by regulation of EE differentiation factors Prospero, Scute and Phyllopod

Our data thus far support a model in which Cph impairs ISC proliferation and EE differentiation, as it is highly induced in the context of ISC/EE-progenitor tumours seen upon either Notch or *tramtrack* RNAi knockdown. Recent years have seen an increased interest in understanding the differentiation process of entero-endocrine cells, which orchestrate metabolism, satiety, immunity and other important organismal responses by the secretion of peptide hormones (Gribble and Reimann, 2019; Guo et al., 2022a; Worthington et al., 2018). In both the *Drosophila* and mammalian intestine, a key role in early EE differentiation is played by members of the Achaete/Scute pro-neural transcription factors (Chen et al., 2018; Schuijers et al., 2015; van der Flier et al., 2009). In addition, the EE-specific TF Pros plays a key role in both the establishment and maintenance of EE cell fate in *Drosophila* (Guo et al., 2024, 2022b). As there is much similarity between the EE-progenitor transcriptional state seen in *ttk^RNAi^* and *Notch^RNAi^* tumours, where Cph is highly expressed, and that in NSCs, we explored previously published DamID DNA binding data on Cph in NSCs (Fox et al., 2022; Tang et al., 2022). The DamID technique as well as its variations NanoDam and TaDa utilise the *E. coli* Dam methylase which, when fused to a gene of interest, will methylate the adenine in GATC sequences in the vicinity of their genomic DNA binding sites for the Dam-TF fusion protein (Steensel and Henikoff, 2000). Regions enriched in these methylated sequences will correspond to the DNA binding intervals of the protein of interest. Methylation peaks for Cph-NanoDam and Cph-TaDa in NSCs identified Cph binding peaks in 5’ promoter regions of several key EE and ISC determinants such as *pros*, *ttk*, *phyl, E(spl)m8-HLH* and *sc* (Figure 6 b, d-g). Furthermore, other genes with binding peaks are key cell cycle regulators such as E2f1 (Figure 6 a) and Cyclin E (data not shown). Furthermore, we identify binding peaks in several genes that are involved in NSC fate and division such as Sequoia (*seq*, data not shown), Miranda as well as Nerfin-1 and the Yorkie co-factor Scalloped that function as downstream transcriptional effectors of the Hippo signalling pathway (Figure 6c, h and data now shown). Nerfin-1 has no described role in ISCs, but in NSC it can physically interact with the Yorkie co-factor Scalloped to maintain neuronal fate. Loss of Nerfin-1 and Scalloped leads to de-differentiation on neurons into a stem cell-like fate (Vissers et al., 2018). Furthermore, Hippo signalling and its downstream transcriptional effector duo Yki/Sd have well-described roles in the regulation of ISC-division (Karpowicz et al., 2010; Ren et al., 2010; Shaw et al., 2010).

**Figure 6.**
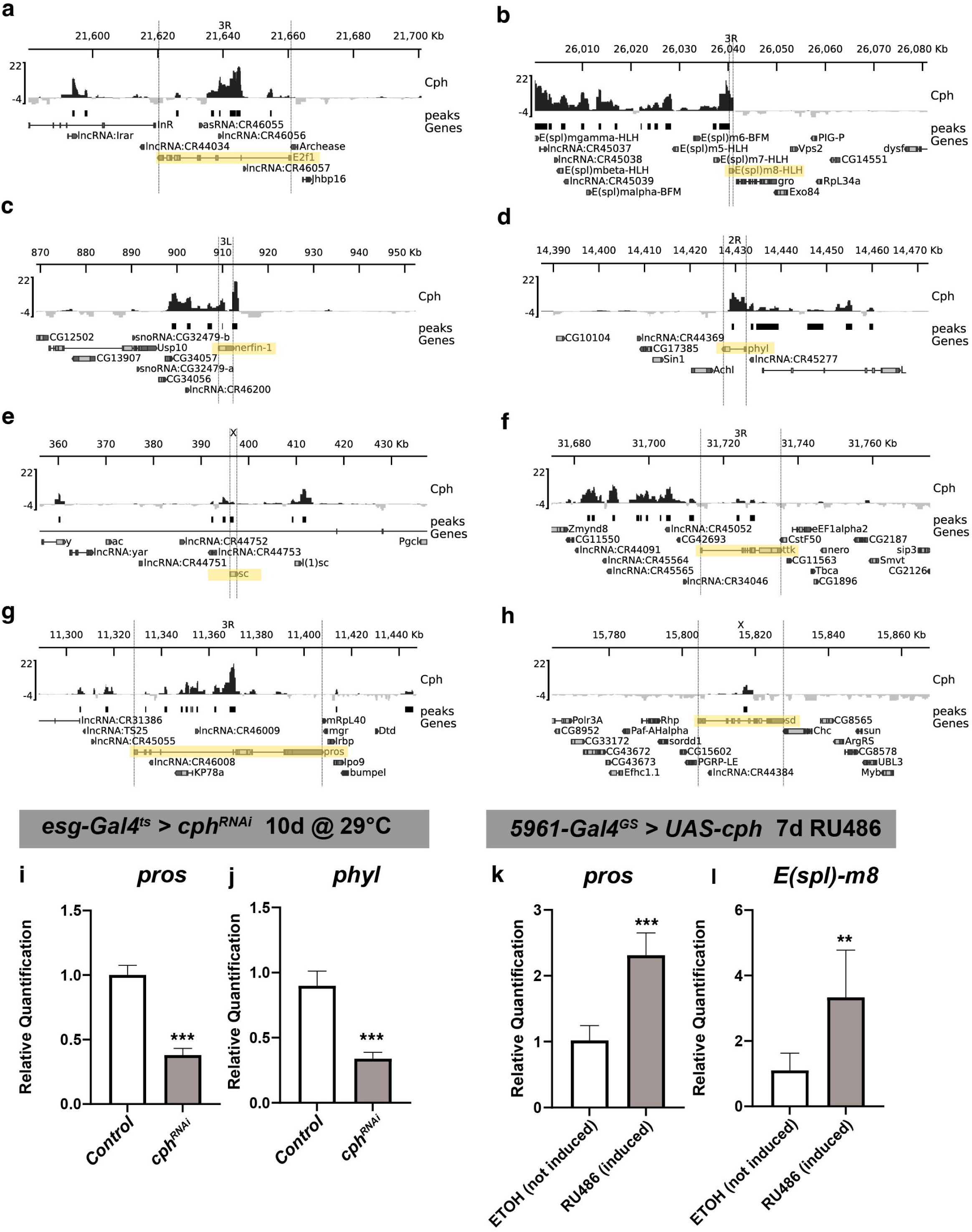
Cph directly regulates key EE-identity factors Prospero, Scute and Phyllopod. **a-h** *cph* Nano-Dam *and* TaDa DamID combined peaks (see Methods) from NSCs from Tang et al., 2022. Cph has binding peaks in the 5’-region of bind *E(spl)m8-HLH, phyl, pros, sc* and *ttk*. Arrows show 5’-3’ direction. **h-i** Cph expression was downregulated by using the *esg-Gal4^ts^ > UAS-mCherry* driver crossed to *UAS-cph^RNAi^* and compared with the *esg-Gal4^ts^ > UAS-mCherry* driver control. RNAi expression was induced for 10 days. **i** *pros* expression was found to be significantly decreased (p=2.9e-4) in *Cph*-depleted samples compared to control. **j** Levels of *phyl* were significantly decreased (*p*= *p*= 1.4e-3). Data is based on two biological repeats and two technical repeats. **k-l** *5961-Gal4^GS^* GeneSwitch induction of Cph induces expression of *pros* and *E(Spl)-m8*. RNA was isolated from induced (RU486) and non-induced (80% EtOH) *5961-Gal4^GS^> UAS-Cph* and used for qPCR with Taqman probes for *pros* and *E(Spl)-m8* genes. **k** *pros mRNA* is significantly (*p*= 0.000488439) upregulated when Cph is overexpressed. Data is based on three technical repeats and compiled from two biological repeats (*n*=15 guts per sample). **l** *E(Spl)-m8* is significantly (*p*= 0.0052233) upregulated when Cph is overexpressed.

Pros is key for both the formation and maintenance of EE cells and is regulated by the proneural gene Scute. The transcriptional regulator *ttk* and its destruction complex consisting of *sina* and *phyl* are part of the same pathway that ultimately regulates the level of *sc* expression. E(spl)m8-HLH is part of the Enhancer of Split ((*E(spl*)) complex, a group of Notch target genes in EBs, which is normally repressed by the activity of Hairless in ISCs to ensure their maintenance (Bardin et al., 2010). The balance between Scute and E(Spl)m8-HLH is necessary for maintaining the balance between ISC quiescence and EE-progenitor formation (Chen et al., 2018; Puig-Barbe et al., 2023). To further analyse whether Cph can directly regulate expression of these genes we examined the expression of *pros* and *phyllopod* mRNA in *esg-Gal4^TS^>cph^RNAi^*guts induced at 29°C for 10 days. Knockdown of *cph* in ISCs/EBs with this driver led to a significant reduction in both *pros* and *phyl* mRNA compared to control (*esg-Gal4^TS^*) (Figure 6 i-j, Supplementary Figure 6).

Next we overexpressed Cph using the GeneSwitch system (Osterwalder et al., 2001), which uses drug-inducible expression through a fusion of the Gal4 DNA binding domain with the trans-activation domain of p65 and the hormone-binding domain of the progesterone receptor. The latter, in the absence of the drug, prevents the fusion protein from entering the nucleus (Han et al., 2000; Osterwalder et al., 2001; Webster et al., 1988). Upon addition of the PR-binding steroid mifepristone (RU-486), the PR-binding domain changes conformation and the fusion protein enters the nucleus, where it can bind UAS sequences and activate transcription. Flies were fed a diet of either 80% EtOH food as a negative control or RU-486 food to activate the system. We employed the *5961-Gal4* GeneSwitch line (*5961-Gal4^GS^*), expressed in ISCs and EBs in the adult midgut, to express *UAS-cph*. Flies were placed on RU-486 food for 7 days to induce Cph expression and guts were isolated for RNA extraction and subsequent qRT-PCR. We observed a significant increase in both *pros* and *E(spl)m8-HLH* mRNA levels in RU486-treated *5961-Gal4^GS^* > *UAS-cph* animals compared to uninduced control (80% EtOH mock-treated) *5961-Gal4^GS^* > *UAS-cph* animals (Figure 6 k-l, Supplementary Figure 6). Hence, Cph binds to and regulates expression of several key regulators that influence ISC-to-EE differentiation.

To further confirm that Cph drives Pros expression and EE differentiation, we overexpressed Cph using *5961-Gal4^GS^* > *UAS-GFP* and quantified the number of Pros-positive cells in the GFP-positive ISC/EB fraction using antibody staining for Pros. We found that at the 7-day timepoint there was a significant increase in the proportion of Pros+ EEs in the GFP+ cells in the *5961-Gal4^GS^* > *UAS-GFP*, *UAS-Cph* genotype compared to the driver control (*5961-Gal4^GS^* > *UAS-GFP*) (Figure 7a-b’’, quantification in 7c). This indicates that an increase in Cph expression leads to increased EE differentiation through its regulation of Pros. In addition, we observed a modest increase in the number of GFP+ cells present within the *5961-Gal4^GS^*> *UAS-cph* guts compared to control (Figure 7d). This further supports a role for Cph in directly driving ISC proliferation.

**Figure 7.**
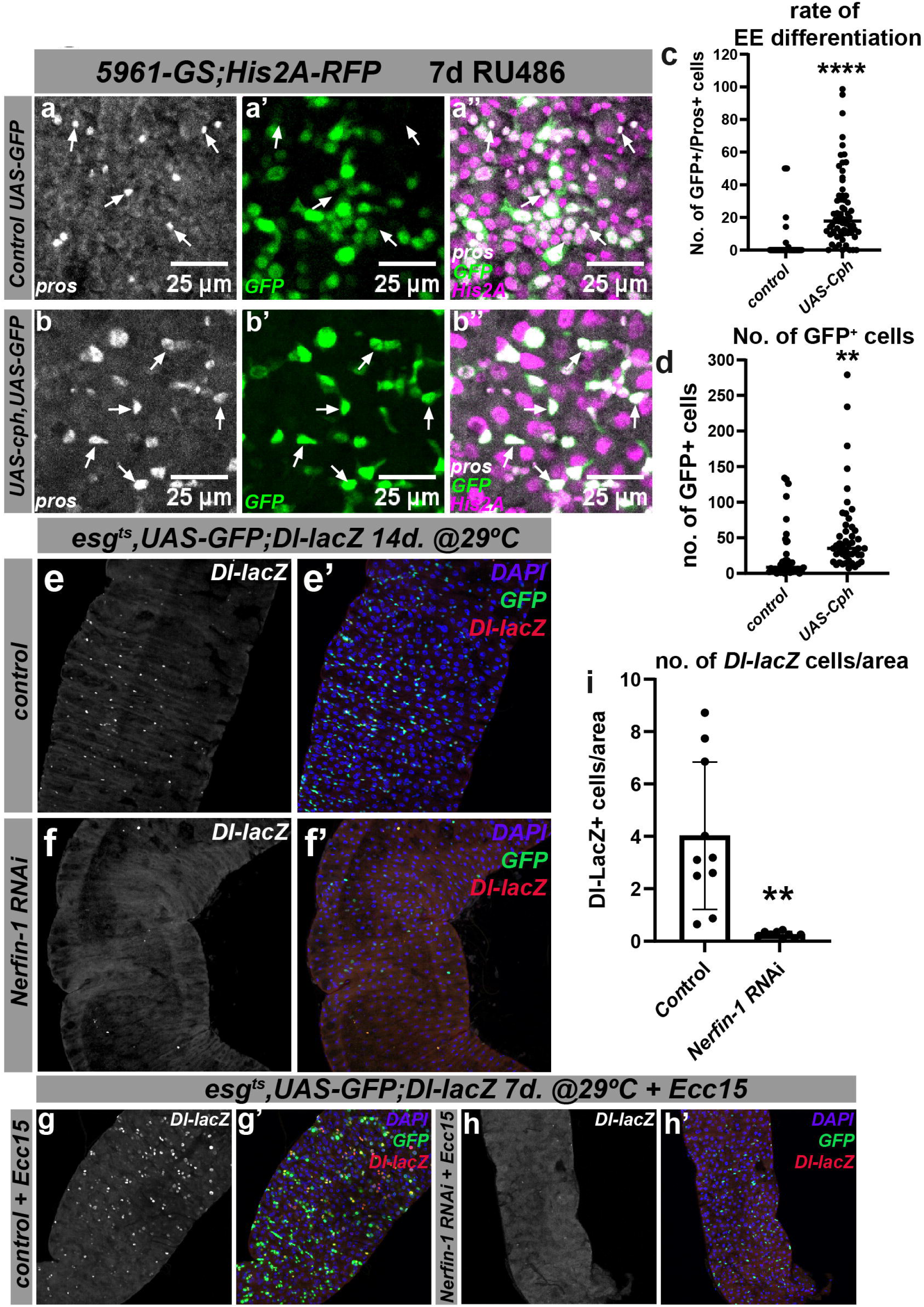
Cph overexpression increases EE differentiation and ISC proliferation through Pros and Nerfin-1. **a-a’’** There were very few Pros^+^ cells within the GFP^+^ cells in the driver control line *5961-Gal4^GS^> UAS-GFP*. Arrows indicate pros^+^ GFP^-^ cells. **b-b’’** A significant (*p*=5.2e-3) increase in Pros^+^/GFP^+^ double-positive cells in the *5961-Gal4^GS^> UAS-GFP*, *UAS-cph* guts (arrows). **c** The rate of EE differentiation as measured by the proportion of GFP^+^ Pros^+^ double-positive cells /Total GFP^+^ cells was significantly increased in guts overexpressing Cph (*p*=2.2e-7). **d** Total number of GFP+ clones were also increased in *5961-Gal4^GS^> UAS-GFP*, *UAS-Cph* overexpressing guts compared to control (*p=*8.7e-3). These results are based on three independent biological repeats with the following cumulative *n*-values: *5961-Gal4^GS^, UAS-GFP;P[His2AV-mRFP] n*=41, *5961-Gal4^GS^, UAS-GFP; UAS-cph/P[His2AV-mRFP] n*=73. **e-f** Images of the *esg-Gal4^ts^, UAS-GFP;Dl-lacZ* line in the driver control (**e-e’**) and *nerfin-1^RNAi^* (**f-f’**) conditions after 14 days induction at 29°C. Note the absence of both GFP^+^ Esg-cells and *Dl-lacZ*^+^ ISCs in the *nerfin-1^RNAi^* condition. **g-h** Impaired regeneration upon *nerfin-1^RNAi^*knockdown after *Ecc15* infection of the *esg-Gal4ts, UAS-GFP;Dl-lacZ* line. Driver control (**g-g’**) and *nerfin-1^RNAi^* (**h-h’**) conditions. **i.** Quantification of no. *of Dl-lacZ*+ ISCs/area of genotypes in (**e-f**) after 14 days induction at 29°C.

To test if Nerfin-1 has a role in ISC proliferation and EE differentiation, we performed knockdown of Nerfin-1 in ISC/EB cells. Levels of Nerfin-1 are almost undetectable in both scRNA-seq atlases for the intestine (Hung et al., 2020; Li et al., 2022). However, knockdown of Nerfin-1 has a dramatic effect on ISC maintenance and proliferation. *nerfin-1* knockdown in *esg-Gal4^ts^, UAS-GFP;Dl-lacZ* animals for 14 days shows widespread loss of Delta+ ISCs and GFP+ ISC/EB nests (Figure 7 f-f’, compare with 7 e-e’, quantification in Figure 7 i). Furthermore, ISC/EBs depleted of *nerfin-1* were not able to respond to intestinal damage by Ecc15 infection, with no visible expansion of the Delta-positive ISC-pool as seen in control animals upon *Ecc15* infection for 24 hours (Figure 7 g-h’). Finally, qRT-PCR showed that guts with Cph overexpression (*5961-Gal4^GS^*> *UAS-cph*) and Notch knockdown both resulted in a significant increase in *nerfin-1* mRNA levels (Supplementary Figure 7). Altogether, our data support a model where Nerfin-1 controls ISC maintenance and proliferation in the adult midgut downstream of Cph.

Based on these results, we propose that Cph controls EE differentiation through direct regulation of key EE determinants such as Pros and Phyllopod. In addition, Cph also regulates genes that control both ISC proliferation and EE-progenitor specification, namely E(Spl)-m8, Scute and the TF Nerfin-1. Our data suggest that Cph promotes the EE fate by directly regulating differentiation factors that act at multiple levels, such as Pros and its upstream regulators Ttk and Phyl.

## Discussion

ISC division and differentiation is mediated by a complex interplay of signalling pathways and transcriptional regulators. Our work has uncovered Cph as a novel key player in the regulation of ISC proliferation and EE differentiation in the adult *Drosophila* intestine. Here, Cph is expressed in both intestinal stem cells (ISCs) and enteroendocrine (EE) cells and is expressed at low levels under homeostatic conditions where ISC proliferation is low. However, Cph expression swiftly increases in contexts where ISC proliferation is high. We have observed this during *Ecc15* infection as well as in the aged intestine, where ISCs proliferate faster due to gut permeability and dysbiosis (Biteau et al., 2010, 2008; Buchon et al., 2009; Rera et al., 2012), but is also characterised by an increase in EE cell differentiation (Choi et al., 2008; He et al., 2018; Tauc et al., 2021).

An increase in EE cells has been found to contribute to the dysregulation of the intestine with age and increased ISC division in both flies and mammals (Amcheslavsky et al., 2014; Tauc et al., 2021; Worthington et al., 2018). The increased Cph-activity that was observed with age correlates with the age-related dysregulation of PcG/Trx gene regulation (Tauc et al., 2021). This age-related epigenetic shift in Polycomb (PcG)/Trithorax (TrxG) gene regulation leads to more EE differentiation with age due to more PcG activity. Interestingly, PcG genes show up as top hits when analyzing the Cph DamID target gene dataset with iCisTarget: a method that identifies regulatory signatures in a set of DNA-binding peaks (Herrmann et al., 2012). Hence, this suggests that Cph might exert its effect by affecting PcG/TrxG-mediated gene regulation. Further experiments will be needed to establish a causal link between Cph and PcG/TrxG gene regulation.

Notch signalling is essential for the control of ISC proliferation and the EC/EE differentiation decision (Micchelli and Perrimon, 2005; Ohlstein and Spradling, 2005). Downstream of Notch in the EB progenitor the EC cell fate regulator Klumpfuss (Klu) acts to ensure lineage differentiation from EB to EC by repressing genes involved in ISC proliferation and EE differentiation. Our previously published Dam-ID data has shown that Klu binds directly to *Cph* and negatively regulates its expression (Korzelius et al., 2019). We find here that *cph* expression is dramatically increased in *Notch-RNAi* induced tumours. Loss of Notch in ISCs leads to the formation of tumours made of neoplastic ISC-like cells and EE-progenitor-like cells (Patel et al., 2015; Wang et al., 2015). These tumours also lack EC-differentiation. We show that *Notch^RNAi^* -driven tumour growth is dependent on Cph, as its knockdown results in decreased proliferation rates and increased survival of Notch RNAi tumour animals. In addition, we find that Cph binds several key cell cycle regulators such as E2F1 and Cyclin E Figure 6a and data not shown). In addition, Cph can bind and regulate the TF Nerfin-1, which was previously found in NSC to physically interact with and regulate transcriptional activity of the Yorkie co-factor Scalloped that mediates the transcriptional response to the Hippo signalling pathway (P. Guo et al., 2019; Huang et al., 2005; Vissers et al., 2018). Yki/Sd have well-described roles in positively regulating ISC proliferation downstream of Hippo signalling (Karpowicz et al., 2010; Ren et al., 2010; Shaw et al., 2010). However, Nerfin-1 seems to act as a repressor of Yki/Sd-driven cell proliferation in the context of neuroblast/NSC and wing-disc cell proliferation and acts as a positive regulator of NSC differentiation by inducing exit from the NSC state (Froldi et al., 2015; Vissers et al., 2018; Xu et al., 2017). This suggests that Nerfin-1 might act independently of Yki/Sd in positively regulating ISC proliferation. Altogether, our data support a key role for Cph in regulating ISC proliferation through various mechanisms, both under homeostatic conditions and conditions of high ISC-proliferation.

Previous reports have described genetic interactions between Cph and Notch signalling. For instance, Cph positively regulates Notch signalling in cone cell specification in the eye (Shalaby et al., 2009). In parallel to this, high Cph expression was also observed in neighbouring mesenchymal myoblast progenitors in an epithelial wing disc tumour model (Boukhatmi et al., 2020; Herranz et al., 2014). These myoblast progenitor cells are also dependent on Notch signalling for their maintenance and proliferation and to prevent differentiation into mature muscle cells (Gunage et al., 2017). This suggests that Cph plays a general role in regulating stem/progenitor cell identity and proliferation in Notch-dependent stem/progenitor cells such as ISCs and myoblasts.

In accordance with this, In the embryonic neuroblast lineage, Cph promotes identity switching from Pdm-to Castor-expressing cells in the temporal cascade that controls differentiation of neuroblasts into neurons along the VNC of the embryo (Fox et al., 2022). Importantly, in contrast to other TFs involved in temporal switching in this system, Cph positively regulates the expression of the next temporal TF in the sequence, rather than repressing the previous TF in the sequence. This suggests that Cph acts mostly as a transcriptional activator of gene expression in this system. This is in line with our qRT-PCR results in the intestine, that show that Cph positively regulates genes involved in EE differentiation and ISC proliferation such as *pros*, *phyl, nerfin-1* and *E(Spl)-m8*. Ectopic expression of Cph in ISCs/EBs results in Pros expression in these cells, suggesting that Cph can trigger EE differentiation. However, we did not find a difference in EE differentiation in *cph* null MARCM clones. This might be due to the small size of these clones and residual Cph protein present in these clones.

In summary, these data support a role for Cph in both ISC proliferation and EE differentiation. This might seem counter-intuitive, as EE differentiation is accompanied by Pros expression, which triggers cell cycle exit by its ability to directly repress cell-cycle genes such as Cdc25/Stg and Cyclin E (Choksi et al., 2006). However, Pros expression is also seen in dividing EE progenitor cells (Zeng and Hou, 2015; Zielke et al., 2014) and Pros activity is also regulated at the post-transcriptional level through sub-cellular localisation (Lai and Doe, 2014). Furthermore, work in neural precursor cells suggests that a phosphorylation-dependent switch on Pros induces neural precursors to switch from a final Pros-driven division to terminal cell cycle exit (Mar et al., 2022). In line with this, it has been shown that EE differentiation is intimately coupled to a final division of the EE-progenitor cells through regulation of the pro-neural factor Scute, which is a positive regulator of Pros-expression (Chen et al., 2018; Puig-Barbe et al., 2023). Hence, ISC division and EE differentiation are intimately coupled, and our data suggest that Cph plays a key role by directly regulating genes involved in both these processes.

The role of Cph in stem/progenitor fate is conserved in mammals, as its orthologues BCL11A and BCL11B play essential roles in the maintenance and function of several adult stem/progenitor cell types. Besides their well-documented roles in immune cell differentiation, ß-Globin regulation and haematological malignancies such as T-ALL (Avram and Califano, 2014; Esrick et al., 2021; García-Aznar et al., 2024), BCL11A and BCL11B (also known as CTIP1 and CTIP2 in mice) play roles in the maintenance of mammary SCs by repression of basal differentiation (Khaled et al., 2015; Miller et al., 2018). Interestingly, *Bcl11b* was found to be required in the mammalian ISCs in the mouse intestine, but loss of one allele of *Bcl11b* resulted in enhanced tumour growth in *Apc^(min/+)^*mice though the modulation of Wnt signalling (Sakamaki et al., 2015). Although Bcl11b plays a tumour-suppressive role in this system compared to the oncogenic role seen in the fly midgut tumours, it does highlight the conserved importance of these family of transcription factors in regulating stem/progenitor behaviour. The contrasting findings in different tissues might be explained by the various molecular interaction partners that have been reported for the Cph orthologues BCL11A and -B. BCL11B was found to interact with both activating Swi/SNF or BAF-chromatin remodelling complexes as well as the repressive NuRD complex that silence chromatin through histone deacetylation activity (Cismasiu et al., 2005; Harb et al., 2016; Kadoch et al., 2013). It is interesting to note that the NuRD interacting domain is conserved between Cph and BCL11B (Yamaguchi et al., 2023). Hence it would be interesting to explore whether this interaction is maintained and with which of these other complexes Cph physically and genetically interacts in fly ISCs and EEs to regulate gene expression and if there are distinct activating as well as repressive chromatin remodelling complexes that can interact with Cph in the context of ISC maintenance and differentiation.

Next to its role in progenitor cell division and differentiation in *Drosophila*, BCL11A/Cph was identified in an genome-wide association study (GWAS) of type 2 diabetes in mammals (Billings and Florez, 2010; Imamura and Maeda, 2011) and the fly orthologue was found to regulate the secretion of the insulin-like peptide dILP2 in Insulin Producing Cells (IPCs) of the fly brain (Park et al., 2014). This role was conserved in mammals, as BCL11A regulates insulin secretion in primary pancreatic islet cells (Peiris et al., 2018). As Cph is expressed in mature EE cells (expressing *Tk_gut_-Gal4*), it would be interesting to see if Cph is required for hormone secretion in gut EE cells. Although dILPs are not expressed by adult EEs (Hung et al., 2020), gut EEs express various hormones that regulate organismal physiology, including the GLP1-analogous peptide hormone NPF and the AstC/Somatostatin hormone that regulate food intake as well as glucose and fat metabolism (Gao et al., 2024; Kubrak et al., 2022; Malita et al., 2022; Yoshinari et al., 2021). It would be interesting to explore whether the expression of Cph in mature EEs is required for their physiology. One example of this is the EE-determinant Prospero, that is not only required during EE-differentiation, but needs to be actively maintained in mature EEs, lest they lose their differentiated fate and partially return to an ISC-like state (Guo et al., 2024, 2022b).

Altogether, our data identify Cph as a critical regulator of ISC proliferation and EE differentiation that becomes deregulated with increased age and in tumourigenesis. As ageing is accompanied by increased stem cell dysfunction as one of its hallmarks (Brunet et al., 2023; Ermolaeva et al., 2018; López-Otín et al., 2023), defining which transcriptional regulators are affected by age in different stem cell tissues will be a crucial first step towards tissue-bespoke therapies to counter stem cell exhaustion with age.

## Materials and Methods

### Fly strains and husbandry

**<u>Bloomington stock centre</u>**: 28324 (*nerfin-1^RNAi^*) 36748 (*ttk^RNAi^*), 58323 (*cph^RNAi^*), 91412 (*M{RFP[3xP3.PB] w[+mC]=Su(H)GBE-smGdP::V5::nls}ZH-2A w[*] P{y[+t7.7] w[+mC]=mira-His2A.mCherry.HA}su(Hw)attP8; P{w[+mC]=UAS-Stinger}2*), 92978 (*cph^[B32]^, P{ry=neoFRT}19A*). **<u>VDRC</u>**: v104402 (*cph^RNAi^*), v341864 (*cph* gRNA). The *UAS-cph (y,w; M{UAS-CG9650.ORF.3xHA.GW}ZH-86Fb)* was obtained from the ORFeome project (FLYORF F000611). The following lines were gifts: *5961-Gal4^GS^* from H. Jasper (Genentech), *Cph-YFP^CPTI1741^* and *1151-Gal4* (*cph-Gal4)* from S. Bray (University of Cambridge, UK), *esg>REDDM (esg-Gal4, UAS-CD8-GFP, UAS-Cas9.p2, UAS-H2B-RFP, tubGal80ts/CyO* from Tobias Reiff *(HHU Düsseldorf),* Tk_gut_-Gal4 (*w;Tk_gut_-Gal4* on 3rd) from Wei Song (Wuhan University).

**Other stocks:** *w^1118^*, *esg-F/O* (*w;esg-Gal4,tub-Gal80ts,UAS-GFP;UAS-flp, Act>CD2>Gal4/TM6B*), *Dl^05151^-lacZ*, *w; If/CyO;UAS-Notch RNAi/Tm6B*, *esg^ts^* (*w;esg-Gal4,UAS-mCherry/(CyO);tub-Gal80^ts^/Tm3,Sb*), MARCM 19A (*hs-FLP,tub-Gal80,FRT19A;tub-Gal4,UAS-GFP*), FRT19A (*y,w,FRT19A*), *cph[A]P{ry=neoFRT}19A (y[1] w[*] CG9650^[A]^ P{ry[+t7.2]=neoFRT}19A/FM7c, P{w[+mC]=GAL4-Kr.C}DC1, P{w[+mC]=UAS-GFP.S65T}DC5, sn[+]),Cphact-F/O (w;tub-Gal80ts, UAS-GFP/CyO,wg-LacZ; UAS-FLP, act>CD2>Gal4/TM6B)*.

### Immunohistochemistry and microscopy

Midguts were dissected in ice-cold 1X phosphate-buffered saline (PBS) fixed in methanol free 4% paraformaldehyde (PFA) for 30 minutes at room temperature. Samples were then washed in 1x PBS with 0.1% Tween-20 for 5 minutes and permeabilized in 1X PBS with 0.15% TritonX-100 for 15 minutes. Samples were then washed 3x 10 minutes in 1x PBS 0.1% Tween-20 before being incubated for 30 minutes in blocking solution (1x PBS, 0.1% Tween-20, 3% bovine serum albumin). Samples were then incubated with primary antibody diluted in blocking solution at 4°C overnight, after which the samples were washed 3x 20 minutes in 1x PBS 0.1% Tween-20 before being incubated in secondary antibody diluted in block solution for 2 hours at room temperature. Samples were then washed 3x 20 minutes in 1xPBS 0.1% Tween-20 before being mounted in Prolong with DAPI (Invitrogen). For Delta staining, we used the N-heptane fixation method. In short, guts were dissected in ice cold 1X PBS before being transferred to 500µL of 4% PFA:N-Heptane 1:1 mix (250µL of N-Heptane and 250µL of 4% PFA diluted from 16% in 1X PBS). Guts are placed on a nutator for 25 minutes, then washed for 2 minutes with 100% Methanol to remove the N-heptane. The tissue was then re-hydrated re-hydrated by washing for 10 minutes in serially diluted methanol in 1x PBS (90-70-50-25-10% methanol in 1x PBS). Subsequently, samples were re-hydrated, washed in 600µL of 0.15% TritonX-100 for 5 minutes and twice more for 15 minutes each. The samples were then blocked for 30 minutes in blocking solution. At this point the protocol continues as stated in the above protocol for regular Immunohistochemistry. Images were acquired using a Zeiss LSM880 confocal microscope using the Zen Black software package. Images were taken of the R4-R5 posterior midgut region using either 20X or 40X magnification. Image analysis was carried out using Fiji (https://fiji.sc/). The following antibodies were used: rabbit anti-phosphorylated Histone H3 (PH3) (1:500; Cell Signalling Technology Cat# 9701), chicken anti-GFP (1:1000; ThermoFisher A10262), mouse anti-Nubbin/Pdm1 (1:10; DSHB Cat# Nub 2D4O), mouse anti-Prospero (1:50; DSHB Cat# MR1A) and mouse anti-Delta (1:100; DSHB Cat# c594.9b).

### Quantification of *Cph-YFP* fluorescence intensity

Fluorescence intensity was measured for each section of the posterior gut imaged. Images were taken in the R4c-R5 region of the posterior midgut. Background mean grey values were measured on a reference image outside the bounds of the gut. The gut was then outlined using the wand tool and the mean grey value was measured. The values obtained for each gut was then subtracted from an average of the mean grey value readings for the background. This generated the corrected gut fluorescence for that portion of the posterior midgut. Fluorescence = (Mean grey value of ROI)-(average Mean grey value of background) Significance was then calculated using the Students T-test with Welch’s correction. P-value significance values are represented by * symbols according to p<0.05=*, p<0.01=**, p<0.001=***, p<0.0001=****.

### RNA isolation and cDNA synthesis

Total RNA was extracted from 15 adult female guts using the RNeasy® mini Kit (Qiagen) according to manufacturer’s instructions. All materials used were RNAse-free. Guts were dissected in ice cold 1X PBS and samples were eluted in 30 µl of RNAse-free water. RNA concentration and purity was measured on a Nanodrop 1000. RNA was immediately used for cDNA synthesis. RNA samples were subsequently stored at -80°C. The SuperScript™ IV Reverse transcription kit (ThermoFisher) was used to generate cDNA according to manufacturer’s specifications. 200 ng of RNA sample was used as input for cDNA synthesis.

### Quantitative real-time PCR (qRT-PCR)

cDNA samples were diluted 1:5 with nuclease-free water. qPCRs were carried out in triplicates with two biological repeats. qRT-PCR was performed using TaqMan™ Fast Advanced Master Mix for qPCR (Thermo Fisher) according to their user guide. Programs were compiled using the QuantStudio™ 3 Design and Analysis software v1.5.2 and qRT-PCR was performed using the QuantStudio™ 3 system. Gene expression was normalized to *Rpl32* expression and relative quantification was calculated using the 2-ΔΔCT method.

### Ecc15 infection

A 50ml culture of *Ecc15* was grown in sterile LB media at 30°C in a shaking incubator overnight. This culture was then spun down at 4000 rpm for 15 minutes at 4°C. The pellet was resuspended in 5ml 5% sucrose solution. 500µl of this solution was dropped onto a piece of filter paper in an empty vial. 15-20 female flies were then placed into this vial for 16-24 hours at 25°C before dissection. As a control 15-20 female flies are also placed in a vial containing 500µl of 5% sucrose solution on filter paper only and placed at 25°C. As a final control, another 15-20 female flies were flipped onto a fresh food vial, also at 25°C. Flies were dissected and stained as described above.

### Tumour survival Assay

Tumour survival assays were carried out based on guidelines specified in Linford et al. (Linford et al., 2013) In each experiment, a cohort of 20-30 flies, hatched in the same 2-day interval, were collected for each phenotype. Survival was measured from the day the animals were placed at the permissive temperature of 29°C. Flies were flipped onto fresh food vials every second day and survival was counted according to Linford et al., (2013). Survival curves were constructed using the Kaplan Meier approach and statistical analysis was carried out using the OASIS 2 software (Han et al., 2016).

### Tumour incidence and growth analysis

Images of the guts were taken using Zeiss confocal microscope and Zen black software. PH3 cell counts were performed manually at the 20X focus using a clicker. Starting in the R5 posterior midgut images were taken at the 20X focus. Two subsequent images were taken travelling from the R5 into the R4 region. The images were analysed using Fiji ImageJ Software. The area of each analysed section was measured using the wand tool. Tumours were counted if there were >8 esg+ cells in a cluster. Each tumour was outlined manually, and its area was taken. Tumour growth was measured as the proportion of the gut taken up by tumours (total area of tumours in gut/total area of gut analysed x 100). Tumour incidence was measured as whether tumours were present or not and this was taken as (no. of guts with tumours/total no. of guts analysed).

### DamID data analysis

Processed DamID bedgraphs of Cph binding in neural stem cells were downloaded from GSE190210 (Tang et al., 2022). These data consisted of two replicates of TaDa profiling, and two replicates of NanoDam profiling. All replicates were scaled by dividing datasets by their standard deviation. As the correlation of all replicates was high, a single averaged binding track for Cph was generated from all four replicates in order to maximise the signal:noise ratio. The averaged track was then used for peak calling via find_peaks v1.0.4 (Marshall et al., 2016), with parameters of fdr=0.01, min_quant=0.92 and step=0.005. Peaks were associated with genes +/- 1kb of the peak via peaks2genes (Marshall et al., 2016). Genomic profiles were generated using pyGenomeTracks (Cubeñas-Potts et al., 2017).

### Pseudo-time expression analysis

scRNA-seq datasets were obtained from GEO (accession number GSE120537) and https://flycellatlas.org. To correct for variability arising from technical and biological effects, we used the *IntegrateData* function in *Seurat* 5.1.0 (Hao et al., 2021; Stuart et al., 2019). The library *Slingshot* 2.12.0 (Street et al., 2018) was used for the trajectory analysis, which focused on ISC/EB cells as the initial state, pEC and EE cells, resulting in the identification of two distinct trajectories.

## Supporting information

Supplementary Figures 1-7 with legends

## Acknowledgements

We thank Heinrich Jasper and Sina Azami for ideas and discussions at the early stage of this project. We also like to thank Tobias Reiff (HHU Düsseldorf), Ingolf Reim (Marburg), Manfred Frasch, Nicholas Tapon (Crick), Michael Taylor (Cardiff) and Sarah Bray (Cambridge), the VDRC (Vienna) and the Bloomington Drosophila Stock Center for providing fly stocks, the DSHB at University of Iowa for antibodies used and Jack Davis and Ian Brown from the Bio-imaging Facility at the University of Kent.

## Author contributions

Conceptualization: J.K. Methodology: E.K., E.J., N.W., JdN and O.M. Formal analysis: E.K, O.M., JdN and J.K. Investigation: E.K., E.J., J.K., O.M., JdN and J.K. Writing E.K, O.M., JdN and J.K. Supervision: JK; Project administration: J.K. Funding acquisition: J.K.

