## Supplementary Figures 1-7 with legends for "The transcription factor Chronophage/BCL11A/B promotes intestinal stem cell proliferation and endocrine differentiation"

### Supplementary Figure 1

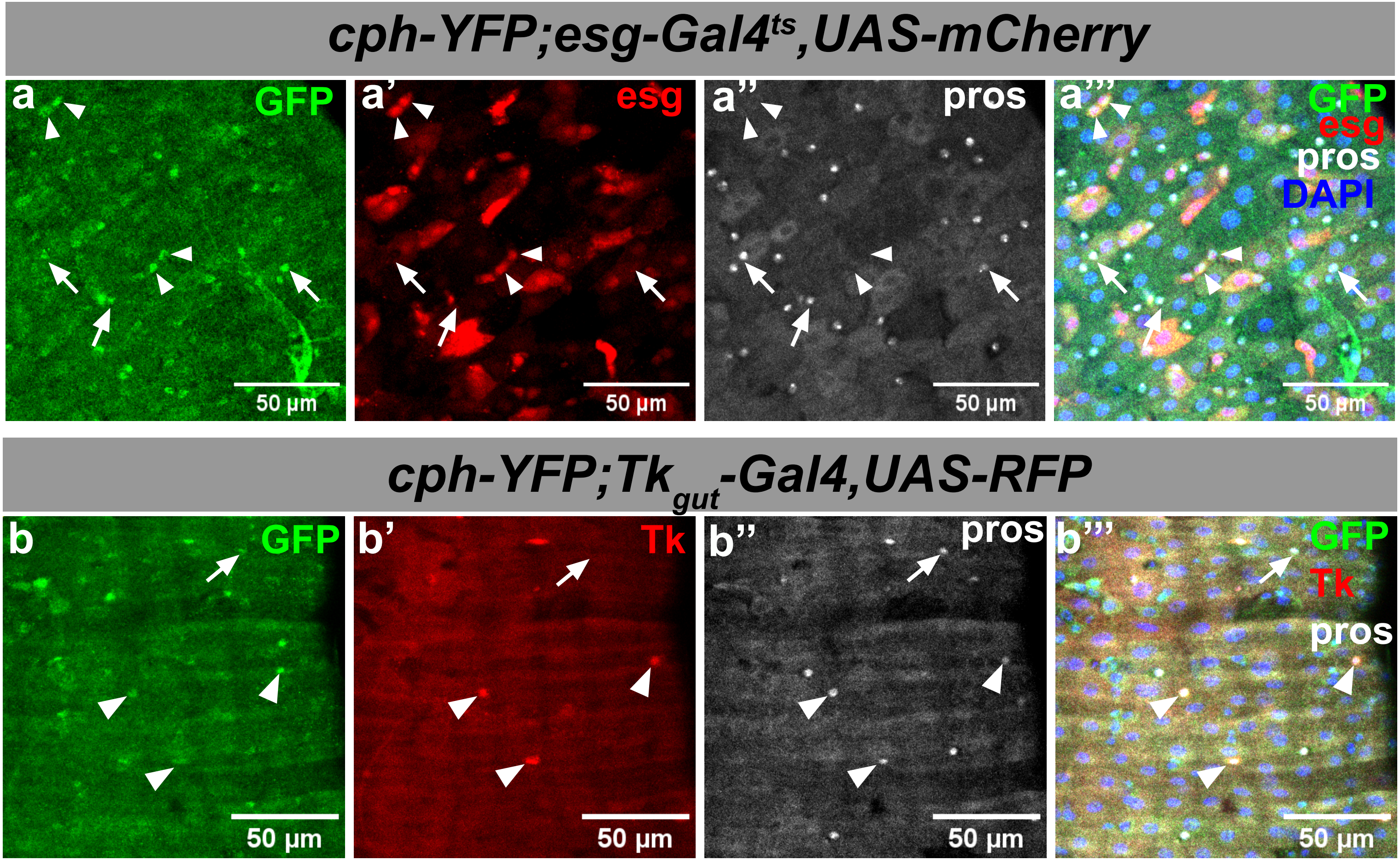

### Supplementary Figure 2

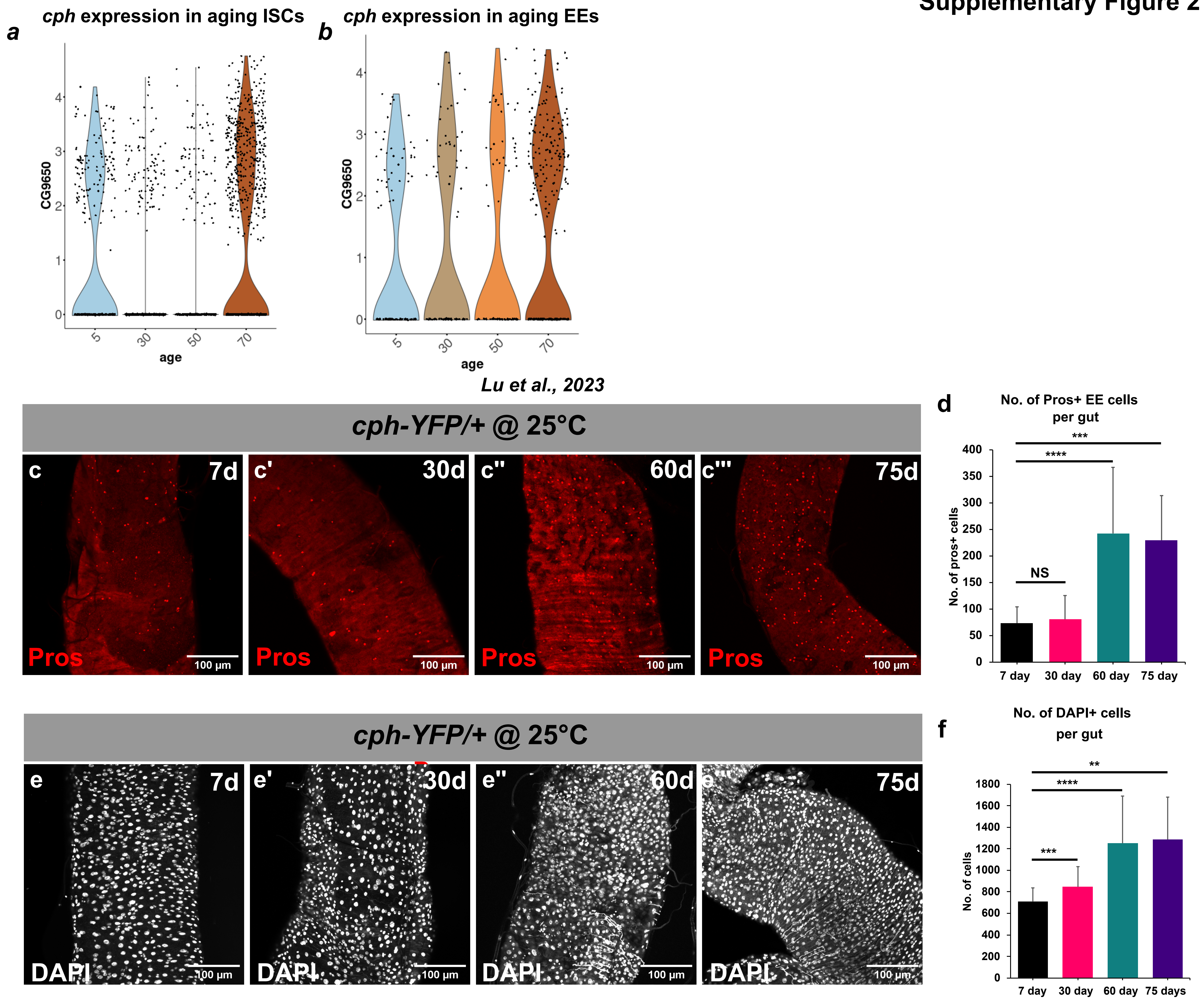

### Supplementary Figure 3

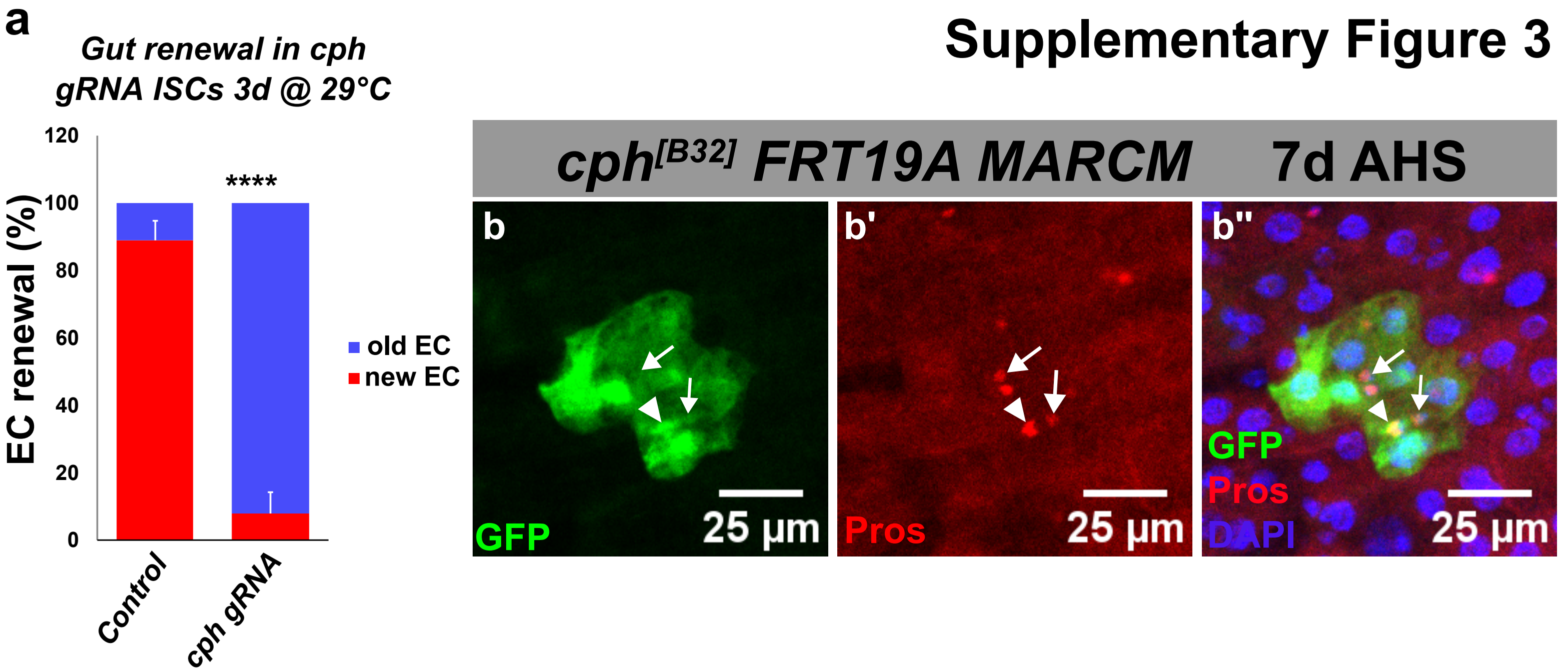

### Supplementary Figure 4

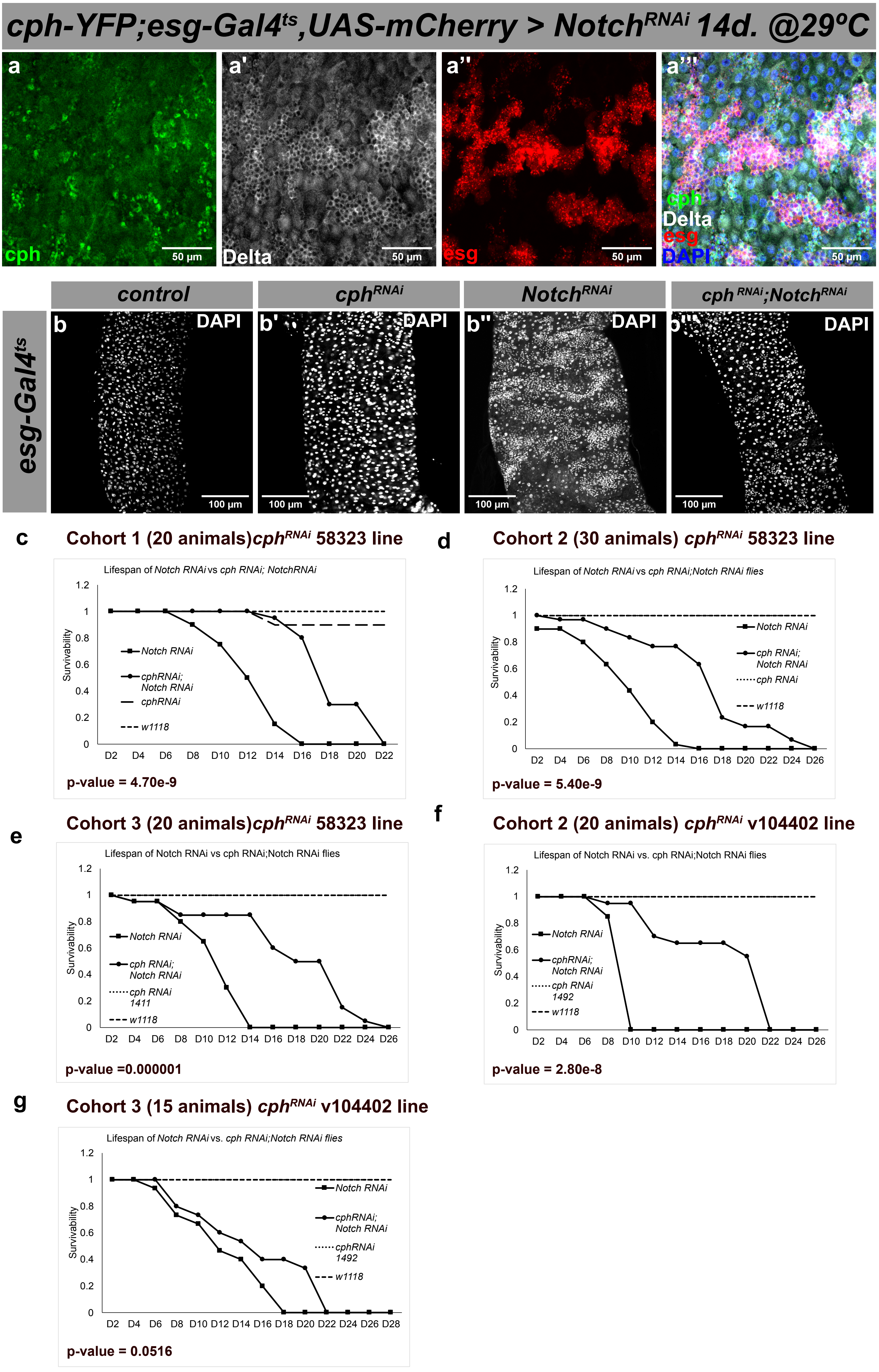

### Supplementary Figure 5

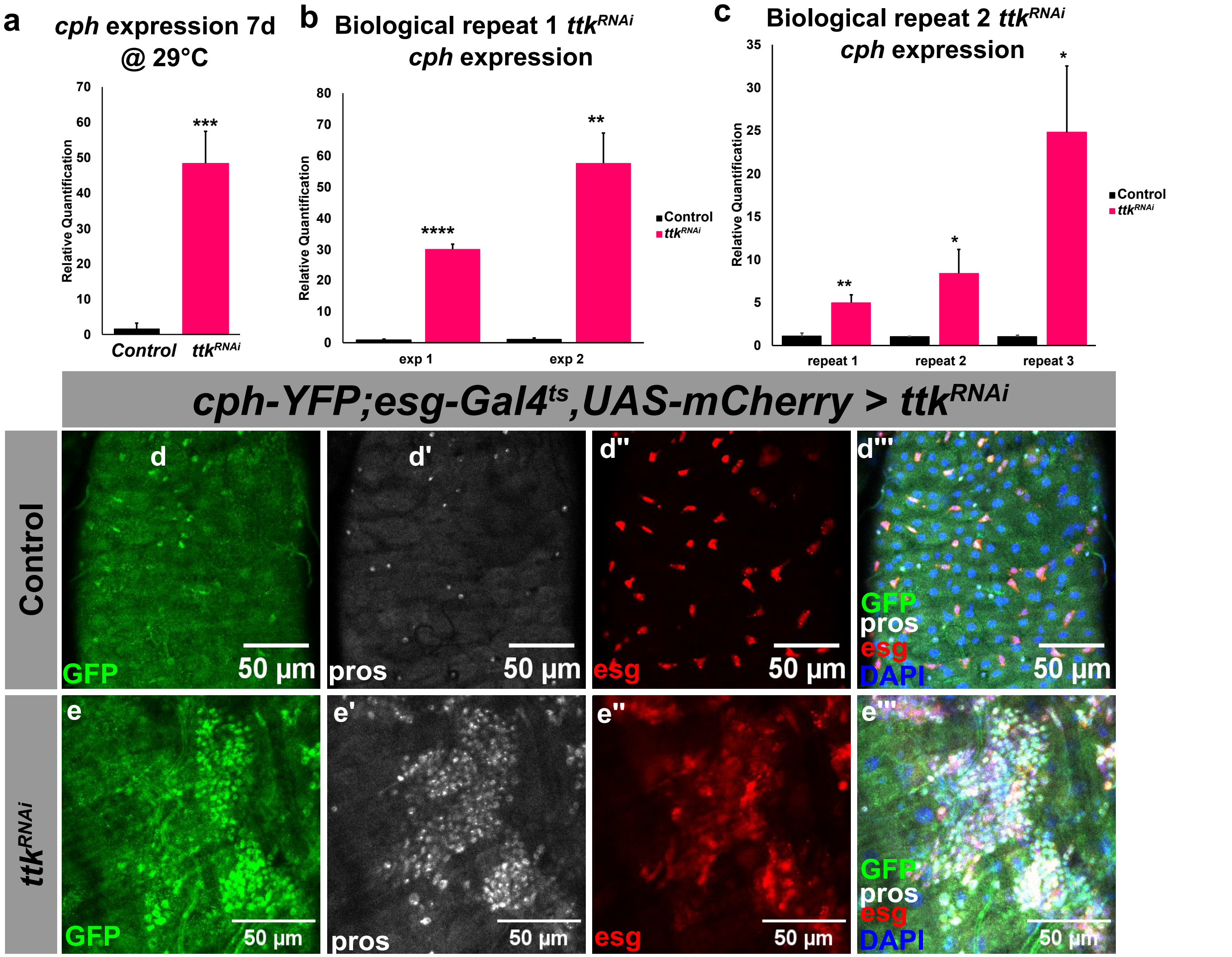

### Supplementary Figure 6

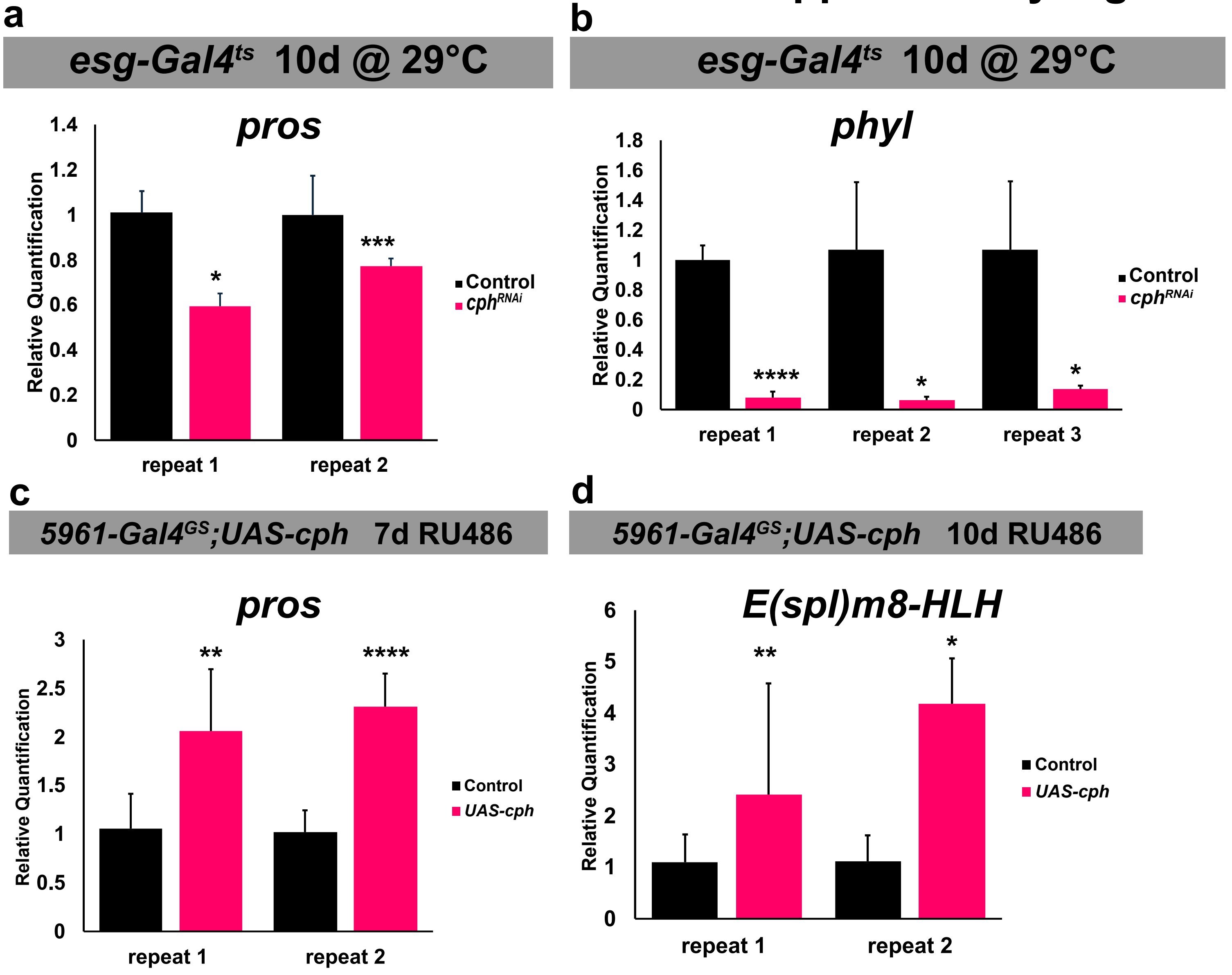

### Supplementary Figure 7

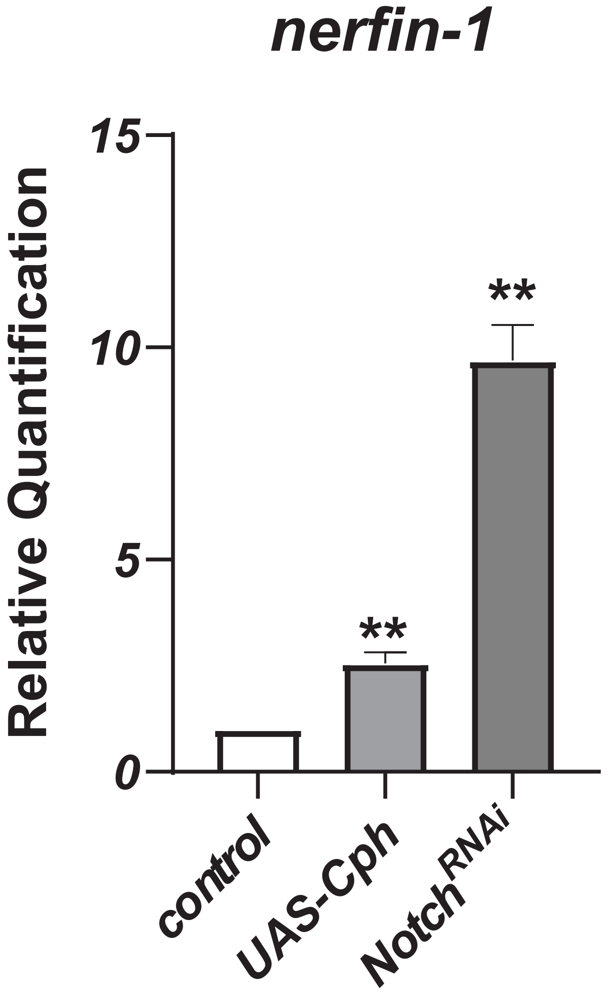
